# Tissue-resident macrophages regulate lymphatic vessel growth and patterning in the developing heart

**DOI:** 10.1101/2020.06.30.179952

**Authors:** Thomas J. Cahill, Xin Sun, Christophe Ravaud, Cristina Villa del Campo, Konstantinos Klaourakis, Irina-Elena Lupu, Allegra M. Lord, Cathy Browne, Sten Eirik W. Jacobsen, David R. Greaves, David G. Jackson, Sally A. Cowley, William James, Robin P. Choudhury, Joaquim Miguel Vieira, Paul R. Riley

**Affiliations:** Burdon-Sanderson Cardiac Science Centre, Department of Physiology, Anatomy and Genetics, University of Oxford, Oxford OX1 3PT, UK.; British Heart Foundation - Oxbridge Centre of Regenerative Medicine, CRM, University of Oxford, Oxford OX1 3PT, UK.; Cardiovascular Development Program, Centro Nacional de Investigaciones Cardiovasculares, CNIC, 28029 Madrid, Spain.; Karolinska Institutet, Department of Medicine Huddinge, Center for Hematology and Regenerative Medicine, Karolinska Institutet, Stockholm SE-14186, Sweden.; Sir William Dunn School of Pathology, University of Oxford, Oxford OX1 3RE, UK.; MRC Human Immunology Unit, Weatherall Institute of Molecular Medicine, John Radcliffe Hospital, University of Oxford, Oxford OX3 9DS, UK.; Division of Cardiovascular Medicine, Radcliffe Department of Medicine, University of Oxford, Oxford OX3 9DU, UK.

**Keywords:** Macrophages, hyaluronan, cell adhesion, cardiac lymphatics, coronaries, vessel growth and patterning

## Abstract

Macrophages are components of the innate immune system with key roles in tissue inflammation and repair. It is now evident that macrophages also support organogenesis, but few studies have characterized their identity, ontogeny and function during heart development. Here, we show that resident macrophages in the subepicardial compartment of the developing heart coincide with the emergence of new lymphatics and interact closely with the nascent lymphatic capillaries. Consequently, global macrophage-deficiency led to extensive vessel disruption with mutant hearts exhibiting shortened and mis-patterned lymphatics. The origin of cardiac macrophages was linked to the yolk sac and fetal liver. Moreover, *Csf1r*^+^ and *Cx3cr1*^+^ myeloid sub-lineages were found to play essential functions in the remodeling of the lymphatic endothelium. Mechanistically, macrophage hyaluronan was found to be required for lymphatic sprouting by mediating direct macrophage-lymphatic endothelial cell interactions. Together, these findings reveal insight into the role of macrophages as indispensable mediators of lymphatic growth during the development of the mammalian cardiac vasculature.

**Summary statement:** Tissue-resident macrophages are indispensable mediators of lymphatic vessel formation during heart development and function to remodel the vascular plexus.

## Introduction

The cardiac vasculature, composed of the coronary circulation and lymphatic vessel network begins to develop from around mid-gestation, at approximately embryonic day (E)11.5 in the mouse embryo. Lymphatic endothelial cells (LECs) expressing the canonical lymphatic prospero homeobox 1 transcription factor (PROX1), vascular endothelial growth factor receptor 3 (VEGFR3) and lymphatic vessel endothelial hyaluronan receptor 1 (LYVE-1), first arise in the vicinity of the sinus venosus (dorsal side) and outflow tract (ventral side) of the murine heart at approximately E12.5 (Klotz et al., 2015). LECs then assemble into a primitive plexus that expands and remodels prenatally in the sub-epicardial surface along the base-to-apex axis and postnatally towards the myocardial layer, to form an extensive lymphatic system that drains lymph from the heart to enable optimal cardiac function (Flaht-Zabost et al., 2014; Klotz et al., 2015). Defects in lymphatic drainage are associated with heart disease, where an increase in tissue fluid content by as little as 2.5% can lead to a 30-40% reduction in cardiac output (Dongaonkar et al., 2010; Laine & Allen, 1991). Conversely, cardiac lymphatics respond to myocardial infarction by re-activating a lymphangiogenic gene expression programme and therapeutic stimulation of this process enhances resolution of macrophage-driven inflammation, promoting tissue repair (Klotz et al., 2015; Vieira et al., 2018). Together, these findings emphasize the importance of the cardiac lymphatic system and the need for a better understanding of the cellular and molecular mechanisms underlying its development.

The ontogeny of LECs integrating within the heart and other organ-based lymphatics has been the focus of a paradigm shift in recent times, with non-venous endothelial precursors now accepted as an additional source of the lineage (Eng et al., 2019; Gancz et al., 2019; Klotz et al., 2015; Martinez-Corral et al., 2015; Stanczuk et al., 2015; Stone & Stainier, 2019; Ulvmar & Makinen, 2016). The precise identity and origin of these non-venous LEC precursors remains somewhat elusive, although genetic lineage tracing experiments have implicated the Tie2/PDGFB-negative transient embryonic hemogenic endothelium of the yolk sac and, more recently, second heart field-derived progenitors as contributing to cardiac lymphangiogenesis (Klotz et al., 2015; Lioux et al., 2020; Maruyama et al., 2019).

Macrophages are myeloid immune cells strategically dispersed throughout the tissues of the body with a vast functional repertoire and emerging plasticity that converges on normal homeostasis, and responses to pathology through mediating inflammation and repair. Macrophages were initially described in sites of physiological cell death within the bulbus cordis of embryonic chick and rat hearts, using light- and electron microscopy (Manasek, 1969; Pexieder, 1975; Sorokin et al., 1994). Subsequently *in vitro* experiments confirmed macrophages as phagocytic cells and, therefore, it was hypothesized that their primary role was to remove debris arising from cell death (Sorokin et al., 1994). Indeed, macrophages are specialised phagocytes with a classical role in engulfing and digesting dying or dead cells, cellular debris and pathogens. Macrophages are also responsible for cytokine production, and act as a source of pro (lymph-)angiogenic factors, such as VEGF-A, VEGF-C and VEGF-D which in turn support tumour growth, vascularisation and dissemination (Qian & Pollard, 2010). In addition to these classical roles in postnatal settings, macrophages have been implicated more recently in organogenesis during embryonic development and tissue regeneration after injury (Aurora et al., 2014; Theret et al., 2019), as well as in the maintenance of arterial tone through regulation of collagen turnover (Lim et al., 2018). In the embryo, macrophages arise initially from the extra-embryonic, transient yolk sac and subsequently from alternative sources within the embryo proper, including fetal liver-derived hematopoietic stem cells (HSC) (Ginhoux & Guilliams, 2016). Yolk sac-derived macrophages seed most tissue-resident niches, which are maintained through adulthood by self-renewal (e.g. microglia in the brain) or gradually replenished by HSC-derived, blood-borne circulating monocyte intermediates, with both populations having distinct roles in tissue injury responses (Hoeffel et al., 2015; Lavine et al., 2014; Stremmel et al., 2018). In the developing heart, the outer mesothelial layer, the epicardium, has been proposed as an important signalling axis to recruit primitive yolk sac-derived macrophages to the subepicardial space (Stevens et al., 2016). The reported timescale for embryonic macrophage recruitment coincides with major morphological changes taking place in the growing heart, namely chamber septation, endocardial cushion formation and valve remodelling, myocardial growth and maturation, development of the electrical conduction system and formation of the coronary and lymphatic vasculature. As such, tissue-resident macrophages have been considered as potential key contributors to some of these processes *via* their functional roles in engulfing dying cells, releasing soluble cytokines, interacting with or recruiting progenitor cells and their potential to transdifferentiate into alternative cell types. Whilst a functional requirement has been identified for cardiac macrophages in valvular remodelling, normal conduction and coronary development (Hulsmans et al., 2017; Leid et al., 2016; Shigeta et al., 2019), there have been no reported roles during cardiac lymphatic development to-date. Given that macrophages contribute to adult lymphangiogenesis within inflammatory, wound healing and tumor microenvironmental settings (Qian & Pollard, 2010; Ran & Montgomery, 2012), as well as in the developing skin where they define nascent vessel calibre (Gordon et al., 2010), we sought to investigate the role of tissue-resident macrophages in the developing heart. Here, we demonstrate for the first time that macrophages are essential for cardiac lymphatic growth and remodeling. Macrophages colonized the developing heart prior to the initiation of lymphatic expansion, closely associating and interacting with the adventitial surface and leading edges of lymphatic vessels where they promote growth and fusion to ensure an adequate coverage over the subepicardial surface. Global genetic ablation of myeloid cells led to hyperplastic and shortened lymphatic vessels in the heart, which failed to branch properly, and to a mis-patterning of the coronary blood vessels. Both extra- and intra-embryonic hematopoietic sources were found to contribute to the tissue-resident macrophage population, and fate mapping, based on canonical myeloid CSF1R and CX3CR1 markers, identified overlapping lineages that contributed to the remodeling of the nascent lymphatic network. In a co-culture model of human lymphatic endothelial cells with human induced pluripotent stem cell-derived macrophages (hiPSC-macrophages), these cells closely associated with tube-forming lymphatic endothelium where they induced sprouting, replicating our *in vivo* findings. Mechanistically, a direct interaction between LECs and hiPSC-macrophages was found to be dependent on macrophage hyaluronan, a linear glycosaminoglycan composed of a repeating disaccharide unit of D-glucuronic acid and N-acetyl-D-glucosamine, previously implicated in cell motility and adhesion during angiogenesis and leukocyte trafficking through the lymph, respectively (Jackson, 2019; Johnson et al., 2017; Lim et al., 2018; Savani et al., 2001). These findings significantly increase our knowledge of (lymphatic) vascular biology and provide further insight into the plasticity and diversity of tissue-resident macrophage function during development. Mechanistic insight into the cellular interactions controlling lymphatic vessel formation is potentially of more widespread interest in terms of understanding how to therapeutically modulate macrophages and lymphatic growth during heart disease.

## Results

### Tissue-resident macrophages are closely associated with developing cardiac lymphatics

To investigate the earliest timepoint at which macrophages were first detected in the developing heart, we initially analyzed published single-cell RNA sequencing (scRNA-seq) data from whole heart at E9.25 and E10.5 (de Soysa et al., 2019; Hill et al., 2019) (Fig. S1). Whilst no macrophage cluster was detected at E9.25 (de Soysa et al., 2019), graph-based clustering followed by dimensionality reduction using Uniform Manifold Approximation and Projection (UMAP) (Becht et al., 2018) of the E10.5 dataset revealed 18 clusters corresponding to the major cardiac cell types (i.e. cardiomyocytes (CMs), endocardial cell (Endo), mesenchymal cells (Mes), epicardial cells (Epi), and second heart field cardiac progenitor cells (SHF)), which also included resident macrophages (Fig. S1A). The identity of the macrophage cluster was defined by the expression of key myeloid markers including receptors for the chemokine fractalkine (*Cx3cr1*) and cytokine colony-stimulating factor 1 (*Csf1r*) (Fig. S1B). Expression of the chemotactic ligand *Csf1* was detected exclusively in the Epi cluster, supporting the hypothesis that epicardial signaling underlies cardiac colonization and seeding of myeloid cells in the subepicardial compartment (Fig. S1B) (Stevens et al., 2016). To validate these findings, we made use of the *Cx3cr1-GFP* reporter mouse, which expresses enhanced GFP under the control of the endogenous *Cx3cr1* locus (Jung et al., 2000). We found that GFP^+^ myeloid cells were present in the developing heart at E10.5 ahead of the onset of coronary and lymphatic vessel growth, being located on the sinus venosus and outer surface of the adjacent ventricular wall (Fig. S1C). Next, to assess a possible role in the formation of the cardiac lymphatic system, we measured the numbers and spatial distribution of tissue-resident macrophages in heart specimens at E12.5 and beyond (Fig. 1). Specifically, flow cytometric analysis of *Cx3cr1^GFP/+^* fetal hearts at E12.5, E14.5 and E16.5 combined with staining for the canonical macrophage marker EGF-like module-containing mucin-like hormone receptor-like 1 (EMR1; henceforth, known as F4/80) (Hume & Gordon, 1983) revealed a distinct population of live singlet cells expressing both markers (Fig. 1A). Moreover, the GFP^+^F4/80^+^ macrophage count increased gradually from 297 ± 60 at E12.5 to 1018 ± 154 at E14.5 and 1569 ± 175 at E16.5 (E12.5 vs. E14.5, *p* < 0.01; E12.5 vs. E16.5, *p* < 0.0001; E14.5 vs. E16.5, *p* < 0.05; Fig. 1A), suggesting active recruitment and/or proliferation of macrophages throughout the time-course of cardiac development. To further characterize the macrophage population residing in the developing heart of *Cx3cr1^GFP/+^* mice, we performed whole-mount immunofluorescence staining using antibodies against the widely-expressed vascular marker endomucin (EMCN; restricted to capillaries and veins from E15.5 onwards (Brachtendorf et al., 2001)) and the lymphatic marker LYVE-1 that is also expressed in macrophages (Klotz et al., 2015; Lim et al., 2018) (Fig. 1B-M). At E12.5, GFP^+^ macrophages were found proximal to the sinus venosus (Fig. 1B,C) and in the outflow tract (Fig. 1D,E), prior to the onset of coronary and cardiac lymphatic vessel formation (Chen et al., 2014; Klotz et al., 2015). Indeed, only a primitive EMCN+ capillary network and GFP^+^LYVE-1^+^ macrophages, but no LYVE-1^+^ lymphatics, were evident in the subepicardial surface at this stage (Fig. 1B-E). At E14.5, GFP^+^LYVE-1^+^ macrophages were more broadly dispersed in both dorsal and ventral surfaces of the developing heart and, in regions covered by vessels, macrophages were found to be in close proximity to, or in direct contact with, the nascent EMCN^+^ coronary and LYVE-1^+^ lymphatic vasculature (compare Fig. 1F,G with Fig. 1H,I). By E16.5, this spatial pattern of distribution was even more evident with GFP^+^LYVE-1^+^ subepicardial macrophages associating with the contiguous patent coronary and lymphatic networks (dorsal and ventral sides), notably alongside vessel branch junctions, leading ends (i.e. tips) and adventitial surface of vessel walls (Fig. 1J-M). Tissue-resident macrophages have previously been implicated in coronary vessel development and maturation (Leid et al., 2016), but their role in cardiac lymphatic vessel growth and remodeling has remained undetermined. To investigate this further we initially validated the findings in *Cx3cr1^GFP/+^* mice, by investigating the developing cardiac lymphatics in transgenic *hCD68-GFP* mice, which report enhanced GFP in macrophages under the control of the human *CD68* promoter and enhancer sequences (Iqbal et al., 2014), combined with VEGFR3, LYVE-1 and PROX1 markers at E16.5 (Fig. 1N-U). Macrophages were found to be co-localized in the subepicardial space with VEGFR3^+^ lymphatic vessels and intimately associated with vessel branching and the leading edges of vessel sprouts (Fig. 1Q, white arrowheads and orthogonal views). Moreover, macrophages were detected bridging adjacent PROX1^+^ lymphatic tips to promote vessel fusion (Fig. 1U, white arrowheads), analogous to the previously reported cellular chaperone role for tissue-resident macrophages during blood vessel expansion in the developing hindbrain and postnatal retina (Fantin et al., 2010). Taken together, these studies reveal that tissue-resident macrophages colonize the embryonic heart prior to the formation of the main vascular networks and adopt a spatial distribution in close proximity to, and in contact with, the forming lymphatics.

**Fig. 1.**
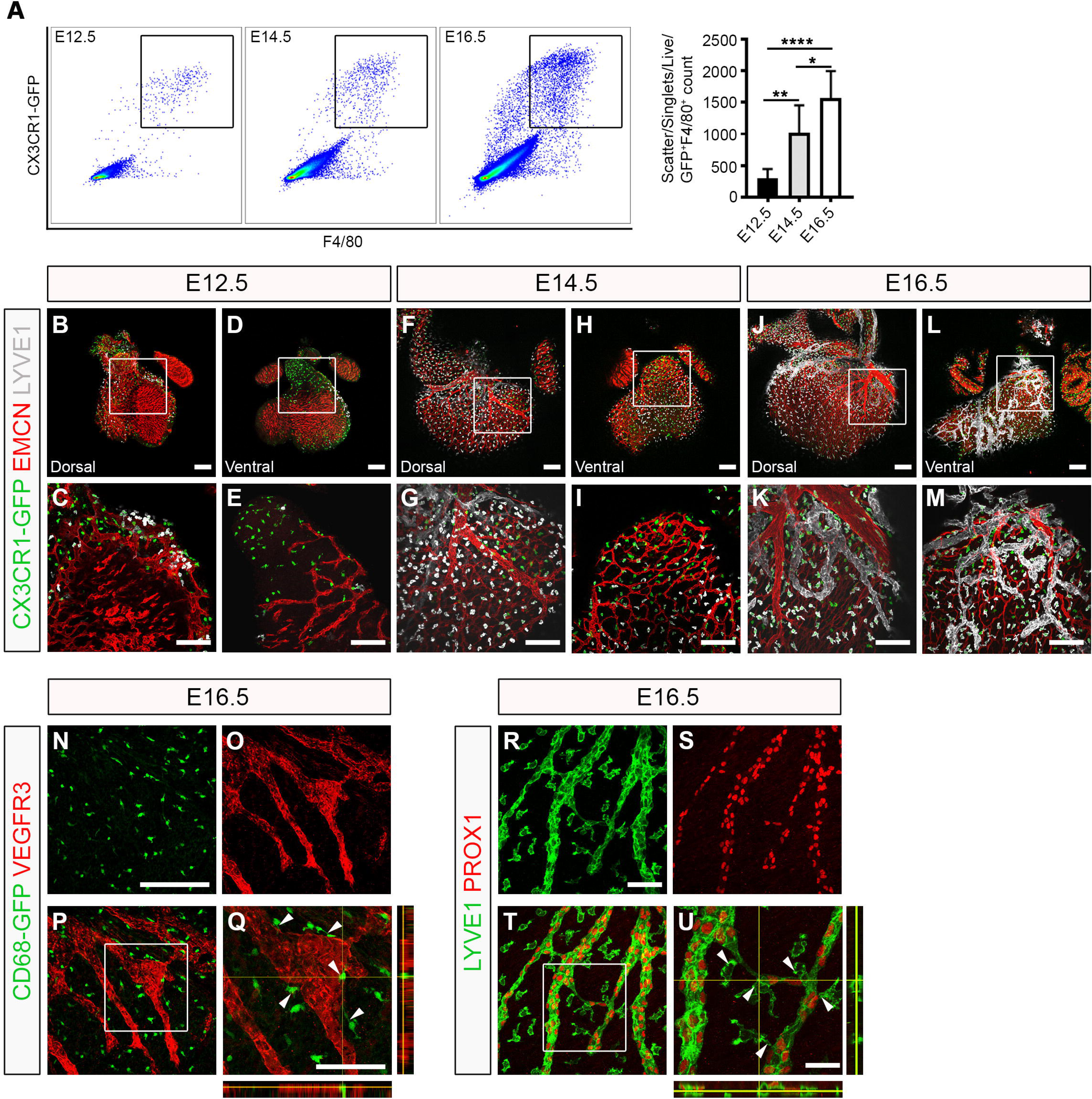
Tissue-resident macrophages are closely associated with the developing cardiac lymphatics. (A) Representative histograms and percentage of tissue-resident macrophages in the developing heart at embryonic days (E)12.5, E14.5 and E16.5 measured by flow cytometry for GFP and F4/80 in *Cx3cr1^GFP/+^* reporter embryos. GFP^+^F4/80^+^ are defined as macrophages. Data represent mean ± SEM; *n* = 6 hearts per group. Significant differences (*p* values) were calculated using one-way ANOVA followed up by the Tukey’s multiple comparison test (\**p* ≤ 0.05; \*\**p* ≤ 0.01; \*\*\*\**p* ≤ 0.0001). (B-M) Whole-mount immunostaining for GFP (green), EMCN (red) and LYVE-1 (white) to visualize the sub-epicardial tissue-resident macrophages, coronary vessels (capillaries and veins) and lymphatic plexus (respectively) in both the dorsal and ventral aspects of hearts-derived from *Cx3cr1^GFP/+^* embryos at E12.5 (B-E), E14.5 (F-I) and E16.5 (J-M). (C,E,G,I,K,M) Magnified views of boxes shown in (B,D,F,H,J,L). Note that LYVE-1 reactivity is detected in the lymphatic endothelium and tissue-resident macrophages. (N-Q) GFP (green) and VEGFR3 (red) immunostaining of whole-mount hearts derived from *CD68-GFP*-expressing embryos at E16.5. (Q) Orthogonal view of the inset box shown in (P); white arrowheads indicate close association of CD68-GFP^+^ macrophages to VEGFR3-expressing lymphatic vessels. (R-U) LYVE-1 (green) and PROX1 (red) immunostaining of whole-mount hearts derived from C57BL6 embryos at E16.5. (U) Orthogonal view of the inset box shown in (T); white arrowheads indicate LYVE-1^+^ macrophages interacting with fusing lymphatic tip cells labelled with PROX1 (nuclear) and LYVE-1 (membrane). All scale bars 100 μm, except in Q 25 μm, R 200 μm and U 50 μm.

### Yolk sac-derived *Csf1r*^+^ and *Cx3cr1*^+^ myeloid lineages associate with lymphatic expansion

To define the origins of macrophages colonizing the developing heart during lymphatic vessel development, we employed genetic lineage tracing using *Csf1r*, *Flt3* and *Cx3cr1*-based mouse models, which in combination capture the main hematopoietic sources from early to mid-gestation, i.e. the transient hemogenic endothelium of the yolk sac and definitive HSCs within the fetal liver as well as the labelling of yolk sac-derived pre-macrophages (Benz et al., 2008; Ginhoux & Guilliams, 2016; Gomez Perdiguero et al., 2015; Kasaai et al., 2017; Stremmel et al., 2018) (Fig. 2). We initially employed the *Csf1r-Mer-iCre-Mer* transgenic model (henceforth referred to as, *Csf1r-CreER* mice) (Qian et al., 2011) crossed with *R26R-tdTomato* reporter mice in which CRE recombinase activity downstream of *Csf1r* enhancer sequences was induced by tamoxifen pulsing at E8.5 (Fig. 2A-D). This phased induction ensured mapping of the spatiotemporal pattern of *Csf1r* activity in the transient yolk sac-derived erythro-myeloid progenitor (EMP) compartment (Gomez Perdiguero et al., 2015; Hoeffel et al., 2015). At E16.5, *Csf1r-CreER^+^* hearts pulsed at E8.5 revealed tdTomato^+^ cells in close proximity to, or in direct contact with PROX1-expressing LECs, with some of these cells exhibiting dual expression of tdTomato and PROX1 (Fig. 2B, white arrowhead). This modest, but reproducible contribution of the *Csf1r*^+^ lineage to lymphatic endothelium (Klotz et al., 2015), supports the hypothesis that a subset of CSF1R^+^ EMPs emerge from the yolk sac at E8.5 to colonize the developing heart and acquire an LEC phenotype, before integrating into the nascent vasculature and contributing to the reported endothelial heterogeneity (Klotz et al., 2015; Lioux et al., 2020; Maruyama et al., 2019). Recently, a yolk sac-derived CSF1R^+^ EMP population was reported to contribute extensively to the endothelium of the nascent vasculature of the developing hindbrain (Plein et al., 2018). However, unlike in the brain, yolk sac-derived CSF1R+ EMPs colonizing the heart appear to give rise largely to tissue-resident macrophages seeding the subepicardial compartment.

**Fig. 2.**
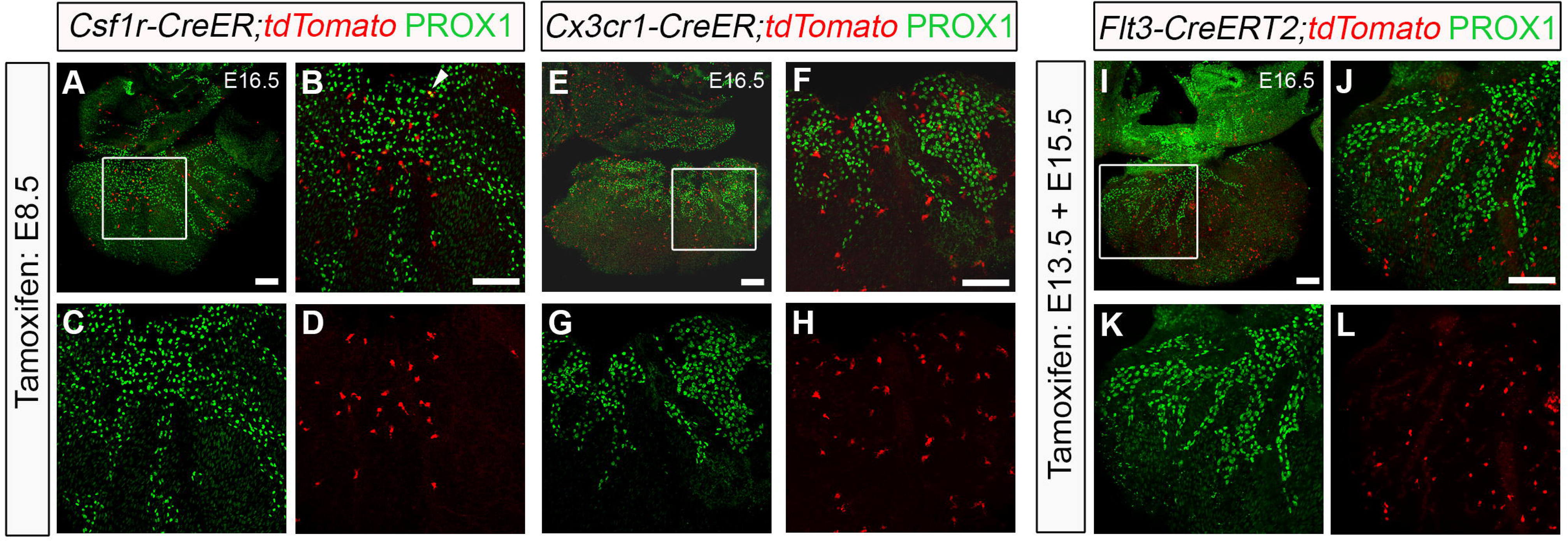
Yolk sac-derived *Csf1r*^+^ and *Cx3cr1^+^* lineages are associated with cardiac lymphatic growth and expansion. (A-D) Genetic lineage-tracing based on the activity of the *Csf1r-CreER;tdTomato* transgene induced by tamoxifen administration at embryonic day (E)8.5. Whole-hearts were analyzed for Tomato (red) and PROX1 (green) expression at E16.5. (B-D) Magnified views of box shown in (A); white arrowhead indicates co-localization of native Tomato and PROX1 immunoreactivity. (E-H) Genetic lineage-tracing based on the activity of the *Cx3cr1-CreER;tdTomato* transgene induced by tamoxifen administration at E8.5. Whole-hearts were analyzed for Tomato (red) and PROX1 (green) expression at E16.5. (F-H) Magnified views of box shown in (E). (I-L) Genetic lineage-tracing based on the activity of the *Flt3-CreERT2;tdTomato* transgene induced by repeated tamoxifen administration at E13.5 and E15.5. Whole-hearts were analyzed for Tomato (red) and PROX1 (green) expression at E16.5. (J-L) Magnified views of box shown in (I). All scale bars 100 μm.

To further characterize the ontogeny of cardiac macrophages, we analyzed the hearts of *Cx3cr1-CreER;tdTomato^+^* embryos, pulsed with tamoxifen at E8.5 (Fig. 2E-H). The *Cx3cr1-CreER* mouse drives expression of the fusion protein CRE-ERT2 (Cre recombinase-estrogen receptor) under the control of the endogenous myeloid-specific *Cx3cr1* locus (Parkhurst et al., 2013) and hence, tamoxifen pulse-chase from E8.5 was anticipated to capture the involvement of yolk sac-derived pre-macrophages (Ginhoux & Guilliams, 2016; Stremmel et al., 2018). Using this approach, we observed widespread contribution from extra-embryonic hematopoietic compartments (i.e. yolk sac) to the macrophage population residing in the developing heart and closely interacting with the nascent PROX1^+^ lymphatic network. *Cx3cr1*^+^ derived cells were exclusively macrophages; we did not observe any direct contribution of LECs within the forming lymphatic plexus (compare Fig. 2E-H with Fig. 2A-D).

To investigate a potential contribution from intra-embryonic definitive HSCs to the tissue-resident macrophage population in the developing heart, we used the *Flt3-CreERT2* mouse model (Benz et al., 2008). Upregulation of FMS-like tyrosine kinase 3 (FLT3; also known as fetal liver kinase 2 or FLK2) is critical for loss of self-renewal of definitive HSCs and *Cre* recombinase-driven by the *Flt3* locus has been demonstrated to label all progeny arising from HSCs (Boyer et al., 2011). At E16.5, *Flt3-CreERT2^+^* hearts pulsed at E13.5 and E15.5 to maximize recombination efficiency resulted in tdTomato^+^ cells residing in close proximity to, or in direct contact with, PROX1^+^ lymphatic vessels on the sub-epicardial surface (Fig. 2I-L), indicating a contribution from the definitive HSC compartment in the fetal liver to the macrophage population residing in heart at later stages of fetal development. Again, as for the *Cx3cr1^+^* lineage, no direct contribution to cardiac lymphatic endothelium was observed.

### Macrophages are required for growth and branching of the cardiac lymphatic network

Our data suggest that yolk sac-derived myeloid lineages, defined by the activity of *Csf1r* and *Cx3cr1* gene expression, migrate to the heart and give rise to macrophages which seed the outer surface of the ventricular wall and interact closely with the neighboring lymphatic vessel network that develops from E12.5 onwards. To determine the functional requirements for these tissue-resident macrophages, we first investigated cardiac lymphatic development in a mouse model deficient for the essential ETS domain-containing transcription factor PU.1 (Fig. 3). *Pu.1* knock-out mice die around birth due to a severely underdeveloped immune system, including a lack of myeloid cells (McKercher et al., 1996; Scott et al., 1994). The number of myeloid cells and macrophages in the developing hearts of *Pu.1*-null mice was determined by flow cytometry for the expression of the pan-leukocyte CD45, myeloid CD11b and macrophage F4/80 markers, and found to be significantly reduced compared to control littermate embryos (CD45^+^CD11b^+^ myeloid cells: 1.35 ± 0.25% vs. 0.02 ± 0.008% of all live cells, *p* < 0.01; CD45^+^CD11b^+^F4/80^+^ macrophages: 85.4 ± 2.77% vs. 0.0 ± 0.0% of all live cells, *p* < 0.0001; Fig. S2A-D). As a consequence, cardiac lymphatic growth and patterning was observed to be severely disrupted in *Pu.1*^-/-^, compared to control littermates at E16.5; with mutant hearts exhibiting hyperplastic and shortened LYVE-1^+^ vessels (vessel length: 12536 ± 1155 μm vs. 7231 ± 734.8 μm; *p* < 0.01), as well as a significant reduction in the number of junctions (i.e. branchpoints; co, 254.2 ± 32.77 vs. *Pu.1^-/-^*, 104.0 ± 10.52; *p* < 0.01) and overall plexus complexity; indicative of a sprouting/branching defect (Fig. 3A-J). A similar phenotype was observed at E19.5, before the onset of embryo demise at perinatal stages (Fig. 3K-T), thus excluding a developmental delay in lymphatic expansion in mutant versus littermate controls. Heart morphology in *Pu.1*-deficient embryos appeared grossly normal, with no evident defects in growth, cardiac septation or compaction of the myocardial layer (Fig. S2E,F), hence ruling out the possibility of a secondary cardiac phenotype contributing to the observed lymphatic defects. However, coronary blood vessel growth and patterning were mildly affected, with null-hearts displaying an overall reduction in subepicardial capillary vessel length and junction number (Fig. 4A-J), as well as exhibiting extra branches of main EMCN^+^-coronary veins (Fig. 4E, white asterisk), compared to control littermates at E16.5. The mis-patterning of EMCN^+^-coronary veins was still evident in *Pu.1*-deficient hearts at E19.5 (Fig. 4K-R, white asterisks in O and Q), whereas the coronary capillary vessel length and junction density were restored to equivalent levels as control littermate hearts (Fig. 4S,T).

**Fig. 3.**
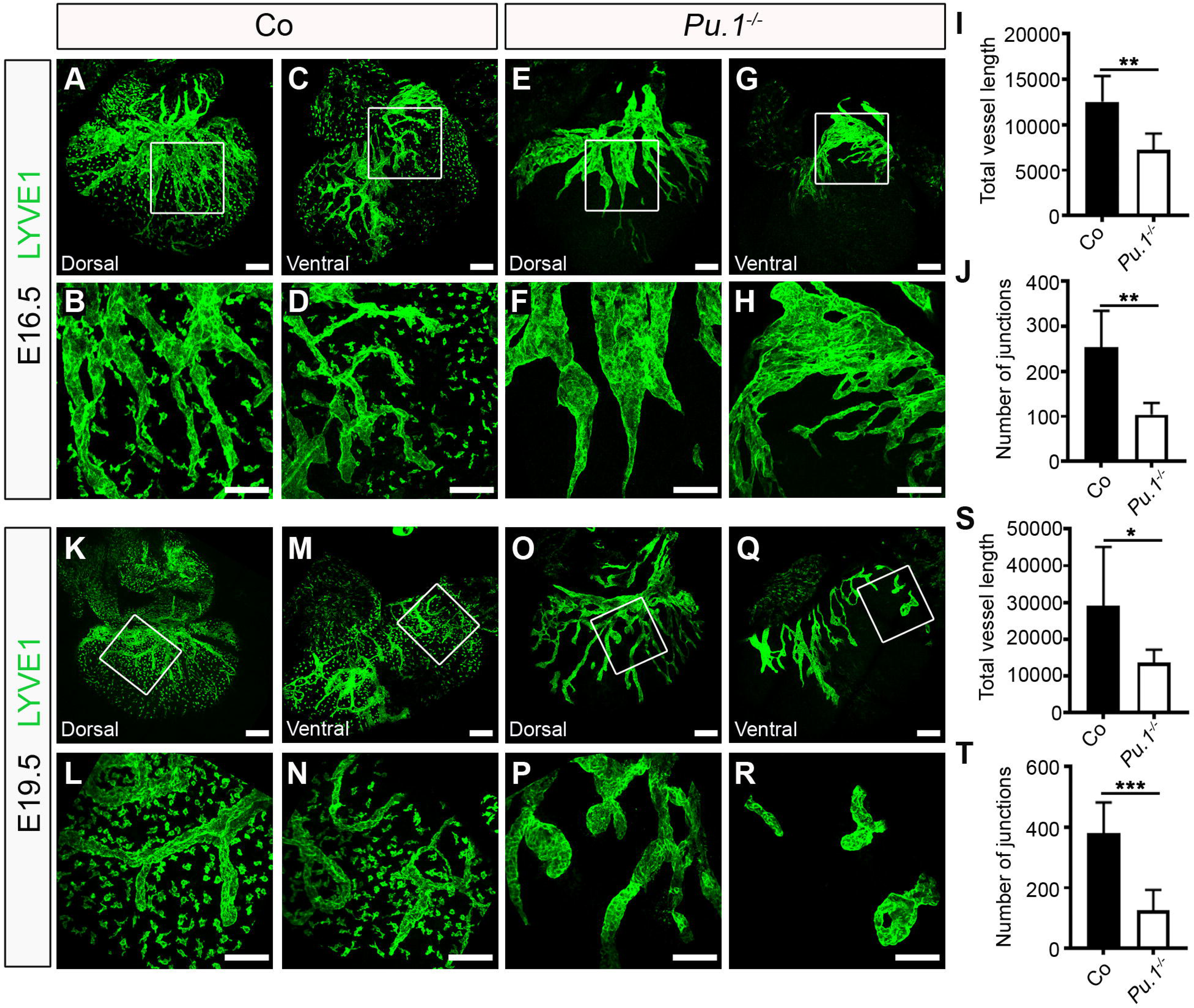
Macrophages are essential for growth and branching of the cardiac lymphatic network. (A-H) Whole-mount immunostaining for LYVE-1 (green) to visualize the sub-epicardial lymphatic plexus in both the dorsal and ventral aspects of hearts-derived from littermate control (co; A-D) or *Pu.1^-/-^* embryos (E-H) at E16.5. (B,D,F,H) Magnified views of inset boxes shown in (A,C,E,G). Note absence of LYVE-1 reactivity in tissue-resident macrophages in *Pu.1^-/-^* hearts, compared to control littermates, confirming absence of macrophages. (I,J) Quantification of total vessel length (μm; I) and number of lymphatic vessel junctions (J) in Co versus *Pu.1^-/-^* hearts at E16.5. Data represent mean ± SEM; *n* = 6 hearts per group. Significant differences (*p* values) were calculated using an unpaired, two-tailed Student’s *t*-test (\*\**p* ≤ 0.01). (K-R) Whole-mount immunostaining for LYVE-1 (green) to visualize the sub-epicardial lymphatic plexus in both the dorsal and ventral aspects of hearts-derived from littermate control (co; K-N) or *Pu.1^-/-^* embryos (O-R) at E19.5. (L,N,P,R) Magnified views of boxes shown in (K,M,O,Q). (S,T) Quantification of total vessel length (μm; S) and number of lymphatic vessel junctions (T) in Co versus *Pu.1^-/-^* hearts at E19.5. Data represent mean ± SEM; Co, *n* = 5 hearts; *Pu.1^-/-^*, *n* = 6 hearts. Significant differences (*p* values) were calculated using an unpaired, two-tailed Student’s *t*-test (\**p* ≤ 0.05; \*\*\**p* ≤ 0.001). All scale bars 100 μm.

**Fig. 4.**
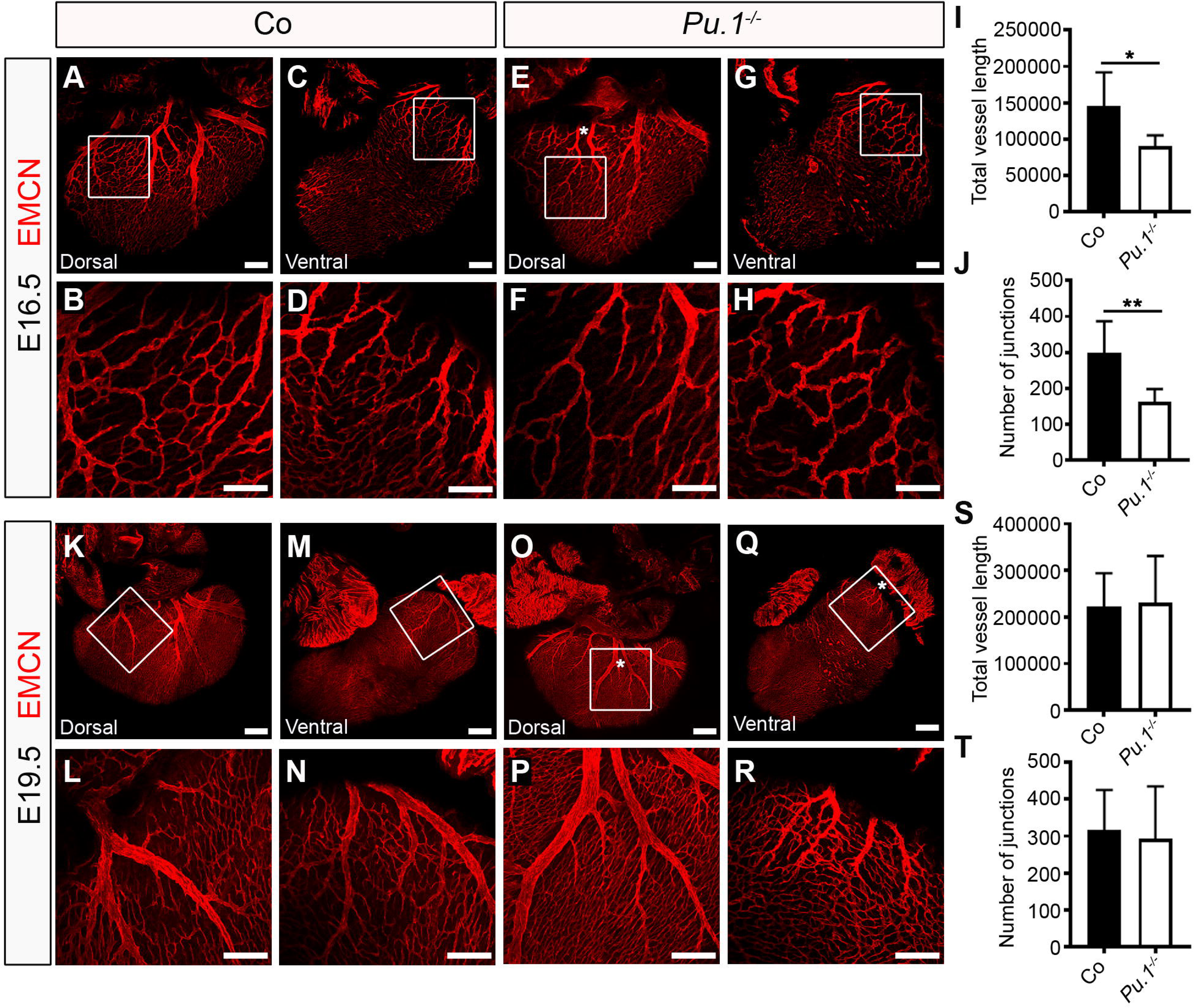
Macrophages regulate coronary growth and patterning. (A-H) Whole-mount immunostaining for EMCN (red) to visualize the sub-epicardial coronary vessels (capillaries and veins) in both the dorsal and ventral aspects of hearts-derived from littermate control (co; A-D) or *Pu.1^-/-^* embryos (E-H) at E16.5. (B,D,F,H) Magnified views of boxes shown in (A,C,E,G). White asterisk indicates patterning defects, i.e. an extra mid-branch, of the coronary veins on the dorsal aspect of *Pu.1^-/-^* hearts (compared E with A). (I,J) Quantification of total vessel length (μm; I) and number of vessel junctions (J) in Co versus *Pu.1^-/-^* hearts at E16.5. Data represent mean ± SEM; *n* = 6 hearts per group. Significant differences (*p* values) were calculated using an unpaired, two-tailed Student’s *t*-test (\**p* ≤ 0.05; \*\**p* ≤ 0.01). (K-R) Whole-mount immunostaining for EMCN (red) to visualize the sub-epicardial coronary vessels (capillaries and veins) in both the dorsal and ventral aspects of hearts derived from littermate control (co; K-N) or *Pu.1^-/-^* embryos (O-R) at E19.5. (L,N,P,R) Magnified views of boxes shown in (K,M,O,Q). White asterisks indicate patterning defects, i.e. extra branches, of the coronary veins on both the dorsal and ventral aspects of *Pu.1^-/-^* hearts (compared O,Q with K,M). (S,T) Quantification of total vessel length (μm; S) and number of vessel junctions (T) in Co versus *Pu.1^-/-^* hearts at E19.5. Data represent mean ± SEM; Co, *n* = 5 hearts; *Pu.1^-/-^*, *n* = 6 hearts. No significant differences were determined using an unpaired, two-tailed Student’s *t*-test. All scale bars 100 μm, except E 1mm.

In a previous study, macrophages in the skin were shown to define dermal vessel caliber by regulating lymphatic endothelial cell proliferation (Gordon et al., 2010). However, we did not observe any gross changes in LEC proliferation in *Pu.1*-null hearts, compared to control littermates at E16.5 (Fig. S2G-N), suggesting inherent differences across organ-specific lymphatic beds. The latter is supported by the emerging recognition of heterogeneity across the lymphatic vascular system and the existence of distinct origins for sub-populations of LECs (Ulvmar & Makinen, 2016).

To confirm a functional role for macrophages in cardiac lymphatic development and to exclude confounding factors arising from the indiscriminate targeting of immune cell development and potentially contributing to the severity of the phenotype in a *Pu.1^-/-^*background, we analyzed cardiac lymphatic development in mice expressing cytotoxic diphtheria toxin A (DTA) under the control of *Csf1r-CreER* and pulsed with tamoxifen at E12.5 (henceforth, *Csf1r-CreER;R26R-DTA*). DTA activation at E12.5, rather than E8.5 was chosen here to enable normal cardiac seeding by yolk sac macrophages. At E16.5, lymphatic vessel growth was disrupted and the number of vessel junctions was significantly reduced, compared to control littermate samples (vessel length: 21493 ± 1050 μm vs. 15940 ± 1507 μm; *p* < 0.05; number of junctions: 237 ± 23.27 vs. 107.0 ± 21.37 μm; *p* < 0.01; Fig. 5A-J). Moreover, a mild phenotype was identified during the development of the EMCN+ coronary vessels, with evident mis-patterning of the main coronary veins (Fig. 5K-T; white asterisk in O), phenocopying the (blood) vascular defects observed in *Pu.1*-deficient hearts (compare Fig. 5O with Fig. 4E). Therefore, ablation of the *Csf1r*^+^ lineage, concurrently with the onset of cardiac lymphatic development, recapitulated some of the vascular phenotypes (lymphatic and blood endothelium) observed in *Pu.1*-null embryos, supporting a functional requirement for yolk sac-derived macrophages in cardiac lymphatic expansion. Differences in phenotypic severity likely reflect a role for other *Pu.1*-dependent immune cells, in addition to macrophages, and/or adverse effects resulting from the disruption of cardiac (embryo proper) seeding by primitive macrophages at early stages of embryonic development. Moreover, it is worth mentioning that the genetic ablation approach used here does not discriminate between *bona fide* tissue-resident macrophage and *Csf1r^+^* EMPs with LEC potential (Fig. 2; (Klotz et al., 2015)). To circumvent this limitation and categorically define a requirement for yolk sac-derived, tissue-resident macrophages, we analyzed the growth of cardiac lymphatics in mice expressing the DTA under the control of *Cx3cr1-CreER* pulsed with tamoxifen at E12.5 (henceforth, *Cx3cr1-CreER;R26R-DTA*). In this model tamoxifen administration of E12.5 leads to selective targeting of yolk sac-derived macrophages, since CX3CR1 is not expressed in fetal liver-derived monocytes or their precursors (Yona et al., 2013). At E16.5, hearts isolated from *Cx3cr1-CreER^+^* embryos pulsed with tamoxifen at E12.5 exhibited comparatively less LYVE1^+^ tissue-resident macrophages in the subepicardial layer, compared to control littermate samples (Fig. 6A-H), and consequently lymphatic vessel growth was found to be disrupted, with vessel length and number of junctions (i.e. vessel branchpoints) significantly reduced (vessel length: 20866 ± 1088 μm vs. 17800 ± 734.8 μm; *p* < 0.05; number of junctions: 240.9 ± 26.02 vs. 168.0 ± 5.542; *p* < 0.05; Fig. 6I,J), akin to *Csf1r-CreER;R26R-DTA* mutant hearts. This phenotype whilst significant was milder than that observed in *Pu.1*-null hearts, which may be explained by the concomitant replacement of yolk sac-derived macrophages by definitive HSC-derived, blood-borne monocyte progenitors throughout the time-course of lymphatic vessel development (Fig. 2I-L) following ablation of the *Cx3cr1^+^* lineage. This compensation by further seeding of tissue-resident macrophages is not a factor in *Pu.1*-mutants which lack the entire myeloid lineage (compare Fig. 3F,H,P,R devoid of any resident macrophages in *Pu.1*-null hearts with Fig. 6F,H where there is evidence of residual or replenished LYVE-1+ macrophages in *Csf1r-CreER;R26R-DTA* mutant hearts). Interestingly, genetic deletion of the *Cx3cr1^+^* lineage also recapitulated the mis-patterning of the main coronary veins observed in *Csf1r-CreER;R26R-DTA* and *Pu.1^-/-^* hearts (Fig. 6K-T; white asterisk in O). Collectively these data suggest a hitherto unidentified function for tissue-resident macrophages in regulating the growth, patterning and expansion of the contiguous cardiac lymphatic system.

**Fig. 5.**
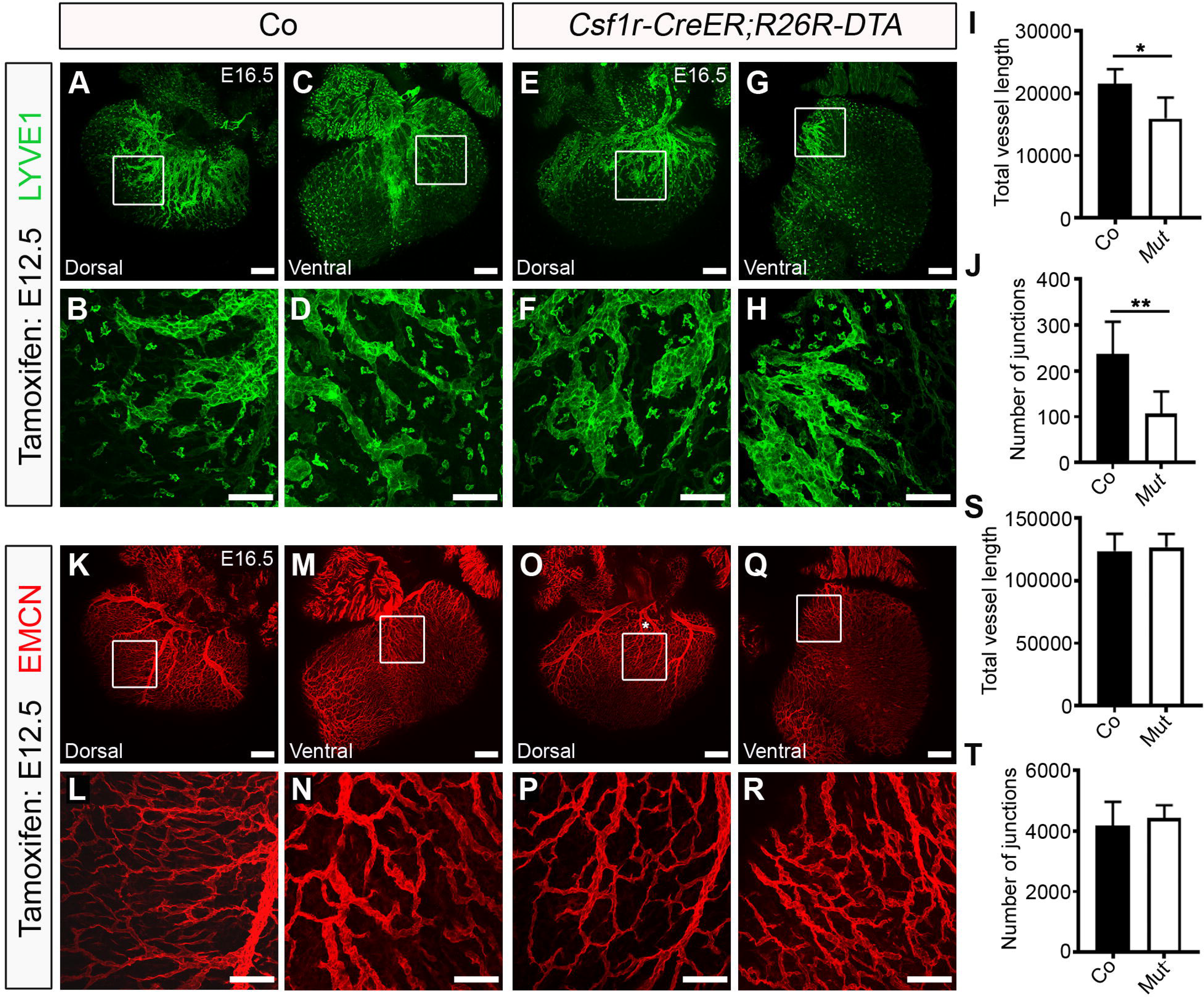
Ablation of the *Csf1r*^+^ lineage impairs cardiac lymphatic growth and branching. (A-H) Whole-mount immunostaining for LYVE-1 (green) to visualize the sub-epicardial lymphatic plexus in both the dorsal and ventral aspects of E16.5 hearts-derived from littermate control (co; A-D) or *Csf1r-CreER;R26R-DTA* embryos (Mut; E-H), tamoxifen-induced at E12.5. (B,D,F,H) Magnified views of boxes shown in (A,C,E,G). (I,J) Quantification of total vessel length (μm; I) and number of lymphatic vessel junctions (J) in Co versus Mut hearts at E16.5. Data represent mean ± SEM; *n* = 5 hearts per group. Significant differences (*p* values) were calculated using an unpaired, two-tailed Student’s *t*-test (\**p* ≤ 0.05; \*\**p* ≤ 0.01). (K-R) Whole-mount immunostaining for EMCN (red) to visualize the sub-epicardial coronary vessels (capillaries and veins) in both the dorsal and ventral aspects of E16.5 hearts-derived from littermate control (co; K-N) or *Csf1r-CreER;R26R-DTA* embryos (Mut; O-R), tamoxifen-induced at E12.5. (L,N,P,R) Magnified views of boxes shown in (K,M,O,Q). White asterisk indicates patterning defects, i.e. an extra mid-branch, of the coronary veins on the dorsal aspect of *Csf1r-CreER;R26R-DTA* hearts (compared O with K), akin to *Pu.1*-null hearts (compared to Fig. 3). (S,T) Quantification of total vessel length (μm; S) and number of vessel junctions (T) in Co versus Mut hearts at E16.5. Data represent mean ± SEM; *n* = 5 hearts per group. No significant differences were determined using an unpaired, two-tailed Student’s *t*-test. All scale bars 100 μm.

**Fig. 6.**
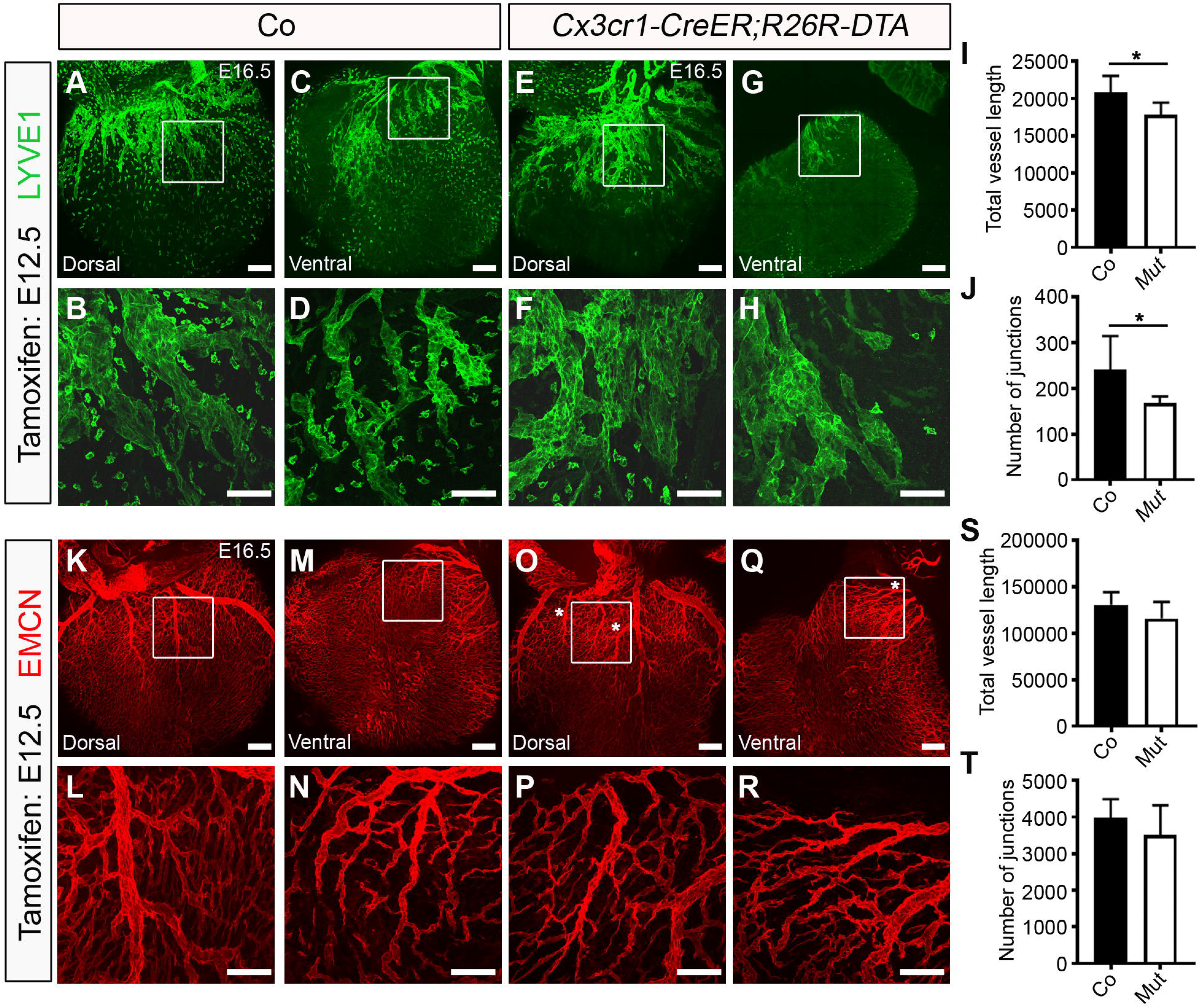
Ablation of the *Cx3cr1*^+^ lineage disrupts cardiac lymphatic remodeling. (A-H) Whole-mount immunostaining for LYVE-1 (green) to visualize the sub-epicardial lymphatic plexus in both the dorsal and ventral aspects of E16.5 hearts-derived from littermate control (co; A-D) or *Cx3cr1-CreER;R26R-DTA* embryos (Mut; E-H), tamoxifen-induced at E12.5. (B,D,F,H) Magnified views of boxes shown in (A,C,E,G). (I,J) Quantification of total vessel length (μm; I) and number of lymphatic vessel junctions (J) in Co versus Mut hearts at E16.5. Data represent mean ± SEM; Co, *n* = 8 hearts; Mut, *n* = 7 hearts; three independent experiments. Significant differences (*p* values) were calculated using an unpaired, two-tailed Student’s *t*-test (\**p* ≤ 0.05). (K-R) Whole-mount immunostaining for EMCN (red) to visualize the sub-epicardial coronary vessels (capillaries and veins) in both the dorsal and ventral aspects of E16.5 hearts-derived from littermate control (co; K-N) or *Cx3cr1-CreER;R26R-DTA* embryos (Mut; O-R), tamoxifen-induced at E12.5. (L,N,P,R) Magnified views of boxes shown in (K,M,O,Q). White asterisks indicate patterning defects, i.e. an extra branch, of the coronary veins on the dorsal and ventral aspects of *Cx3cr1-CreER;R26R-DTA* hearts (compared O,Q with K,M), akin to *Pu.1*-null and *Csf1r-CreER;R26R-DTA* hearts (compared to Figs 3,5). (S,T) Quantification of total vessel length (μm; S) and number of vessel junctions (T) in Co versus Mut hearts at E16.5. Data represent mean ± SEM; Co, *n* = 8 hearts; Mut, *n* = 7 hearts. No significant differences were determined using an unpaired, two-tailed Student’s *t*-test. All scale bars 100 μm.

### Macrophage hyaluronan promotes lymphatic cell network formation and sprouting

The close proximity of tissue-resident macrophages to the developing cardiac lymphatics and evident association with the forming lymphatic endothelium, suggests their role in regulating growth and patterning is most likely mediated by cell-cell contact at the leading edges of fusing branches or the adventitial surface of vessel walls (Figs 1-6). To further explore this possibility, we modeled LEC-macrophage interactions during lymphatic capillary tube formation and sprouting in a human *in vitro* setting, by co-culturing human primary lymphatic endothelial cells and human induced pluripotent stem cell (hiPSC)-derived macrophages-labeled with a red fluorescent protein (RFP) reporter (Fig. 7; Movie S1). The hiPSC-derived RFP^+^ macrophages used here have been previously characterized and shown to exhibit fetal-like properties, mimicking the expression profile and function/activity of tissue-resident macrophages (Buchrieser et al., 2017; Haenseler et al., 2017; van Wilgenburg et al., 2013). In tube formation assays, hiPSC-RFP+ macrophages were found in close proximity to, and associated with LECs, changing cell shape to guide LEC extension and fusion to a neighboring LEC, leading to vessel-like structures (Fig. 7A-F; white arrowheads), analogous to the behavior we observed for embryonic macrophages in the developing heart (Fig. 1). Time-lapse live imaging revealed this to be a highly dynamic process, with individual RFP-labelled cells uncoupling from an individual LEC once a tube was formed and scanning the neighboring micro-environment for further LECs undergoing an equivalent transformation (Movie S1). Likewise, coincident with most of the tube-like structures becoming organized into a plexus, hiPSC-macrophages expressing CD68 were found directly associated with the forming tubes and branching nodes (Fig. 7G-K), phenocopying the behavior observed *in vivo* for tissue-resident embryonic macrophages interacting with cardiac lymphatics (e.g. Fig. 1Q). To model lymphatic sprouting, as an essential first step in lymphatic growth and patterning, we employed two independent three-dimensional (3D) assays based on co-culturing primary human LEC-coated micro-beads (hereafter referred to as the Beads assay) or aggregates of human LECs (the Spheroids assay) with iPSC-derived macrophages (Fig. 7L-X)(Schulz et al., 2012). In the former assay, co-culturing LEC-coated micro-beads with macrophages led to a significant increase in the number of lymphatic-like sprouts per micro-bead, compared to control culture conditions with macrophage:LEC media alone, with RFP-expressing macrophages adjoining and in direct contact with the sprout-leading LEC (control conditions, 100% vs. +macrophages, 173.7 ± 12.90% sprouting activity; *p* < 0.001; Fig. 7L-P). A similar increase in lymphatic sprouting was observed in the spheroids assay (control conditions, 100% vs. +macrophages, 130.9 ± 9.387% sprouting activity; *p* < 0.01; Fig. 7Q-X). To gain insight into the molecular mechanism(s) underpinning stimulation of LEC sprouting by macrophages, we considered paracrine secretion of VEGF-C, a potent lymphangiogenic factor known to be expressed by cardiac macrophages in a myocardial infarction setting (Vieira et al., 2018), and cell adhesion molecules previously implicated in leukocyte-endothelium interactions during immune response, specifically integrin subunit β2 (ITGB2; also known as CD18)- and hyaluronan (HA)-dependent pathways (Gahmberg, 1997; Jackson, 2019; Johnson et al., 2017). Whilst *VEGFC* expression was undetectable, iPS-derived macrophages expressed both *ITGB2* and *ITGAM* (also known as CD11b), the heterodimeric components of the essential macrophage antigen-1 (MAC-1) receptor that binds to endothelial intracellular adhesion molecule-1 (ICAM-1) during leukocyte arrest, rolling and transmigration across the blood vessel wall (Gahmberg, 1997). Notably, they also exhibited a dense coat of HA (previously termed the HA glycocalyx (Johnson et al., 2017; Lawrance et al., 2016)) and expressed the HA-binding receptors *LYVE-1*, *CD44* and *HMMR* (hyaluronan-mediated motility receptor; also known as CD168) (Fig. S3A-D). To discriminate between potential HA and CD18-mediated adhesion to endothelium, we pre-treated iPSC-derived macrophages with hyaluronidase (HAase) to deplete surface glycocalyx-associated HA (Johnson et al., 2017; Lawrance et al., 2016), or with siRNA oligonucleotides targeting *ITGB2/CD18* to specifically knockdown expression of this β integrin subunit without effecting that of the related *ITGAM* expression (Figures S3B-F). We found that loss of macrophage *ITGB2/CD18* had no effect on sprouting activity, but macrophage pre-treatment with HAase significantly impaired LEC sprouting in both bead and spheroid sprouting assays (beads assay: +macrophages, 173.7 ± 12.90 vs. HAase, 96.90 ± 7.205% sprouting activity; *p* < 0.001; spheroids assay: +macrophages, 130.9 ± 9.387 vs. HAase, 80.11 ± 5.588% sprouting activity; *p* < 0.001; Fig. 7O-X), indicating a requirement for macrophage hyaluronan in human lymphatic endothelial growth. Importantly, yolk sac-derived macrophages residing in the HA-rich layer of the epicardium/subepicardium compartment of the developing mouse heart were also found to lack VEGF-C expression and displayed membrane-bound HA (Fig. S4). Taken together with our *Pu.1^-/-^*, *Csf1r-CreER;R26R-DTA* and *Cx3cr1-CreER;R26R-DTA* mutant analyses (Figs 3,5,6), these findings suggest a critical requirement for an evolutionarily-conserved (mouse to human) HA-dependent mechanism underpinning direct cell-cell contact/adhesion between macrophages and LECs during vessel extension and remodeling.

**Fig. 7.**
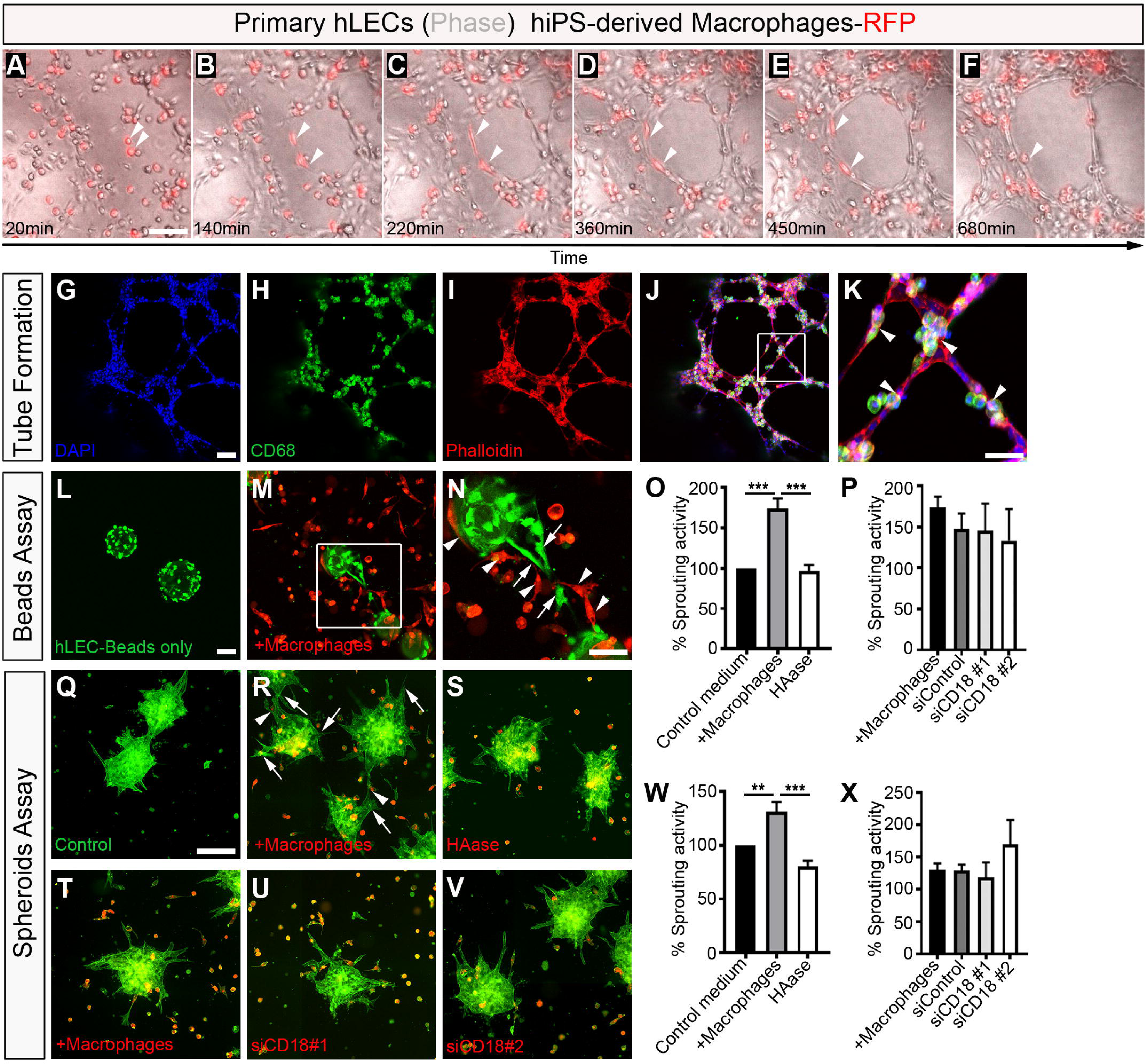
Macrophages promote lymphatic cell plexus formation and sprouting. (A-F) Representative frames of time-lapse experiments using primary human lymphatic endothelial cells (hLECs; phase-contrast) co-cultured with human iPS-derived macrophages labelled with RFP (red), showing close association between macrophages and LECs and a change in macrophage cell morphology (white arrowheads) contacting the expanding lymphatic plexus to facilitate LEC tube formation. (G-K) DAPI (blue), CD68 (green) and Phalloidin (red) staining of primary hLEC and hiPSC-macrophage co-cultures. (K) Magnified view of box shown in (J). (L-N) Representative staining of lymphatic capillary sprouting from beads coated with primary hLECs only (green; control medium; L) or co-cultured with hiPSC-derived macrophages-RFP (red; +macrophages; M). (N) Magnified view of box shown in (M). (O,P) Quantification of % sprouting activity defined as (number capillary sprouts/number of beads) x 100. Data was normalized against the control group. Data represent mean ± SEM; Control, *n* = 6; +Macrophages, *n* = 6; HAase 30U/mL, *n* = 3; siControl, *n* = 3; siCD18 #1, *n* = 3; siCD18 #2, *n* = 3 independent experiments, with three replicates/experiment. Significant differences (*p* values) were calculated using one-way ANOVA followed up by the Tukey’s multiple comparison test (\*\*\**p* ≤ 0.001). (Q-V) Representative staining of lymphatic capillary sprouting from hLECs aggregates/spheroids only (green; control medium; Q) or co-cultured with hiPSC-derived macrophages-RFP (red; +macrophages; R,T), hiPSC-derived macrophages-RFP pre-treated with HAase 30 U/mL (S), or co-cultured with hiPSC-derived macrophages-RFP pre-treated with siRNA oligonucleotides against *ITGB2*/*CD18* (siCD18 #1, U; siCD18 #2, V). (W,X) Quantification of % sprouting activity defined as (number capillary sprouts/number of beads) x 100. Data was normalized against the control group. Data represent mean ± SEM; Control, *n* = 5; +Macrophages, *n* = 4; HAase 30U/mL, *n* = 4; siControl, *n* = 3; siCD18 #1, *n* = 3; siCD18 #2, *n* = 3 independent experiments, with three replicates/ experiment. Significant differences (*p* values) were calculated using one-way ANOVA followed up by the Tukey’s multiple comparison test (\*\**p* ≤ 0.01; \*\*\**p* ≤ 0.001). All scale bars 100 μm, except N 50 μm and Q 250 μm.

## Discussion

Here, we have revealed a novel function for macrophages residing in the developing heart as lymphatic vessel “remodelers” (Figs 1-7). Our findings indicate that primitive macrophages migrate from the yolk sac to the developing heart between E9.25 to E10.5 to initially colonize the outer surface of the ventricular wall, where they reside in the epicardium/subepicardium compartment prior to cardiac lymphatic vessel formation. This compartment has previously been suggested as an essential signaling axis, with epicardial disruption downstream of Wilms’ tumor 1 (*Wt1*) gene deficiency impairing cardiac recruitment of yolk sac macrophages (Stevens et al., 2016). However, the identity of the epicardial signal(s) required for this process has remained elusive. Our scRNA-seq analysis of E10.5 hearts (Fig. S1) revealed that expression of the cytokine *Csf1* was exclusively expressed in epicardial cells. CSF1 regulates the differentiation of most macrophage populations, including primitive yolk sac-derived, and is a potent macrophage-chemoattractant (Ginhoux et al., 2010) and, therefore, is a likely mediator of macrophage recruitment into the subepicardial region of the developing heart. Future studies investigating the control of *Csf1* activity by epicardial factors such as WT1 should provide insight into the molecular basis leading to cardiac seeding by yolk sac-derived macrophages.

The epicardium plays an essential role during heart development by secreting paracrine signals stimulating cardiomyocyte proliferation and maturation, as well as supporting cardiac vasculature expansion (Simoes & Riley, 2018). Within the subepicardial space, tissue-resident macrophages co-exist with both coronary and lymphatic endothelium. Our study attributes novel function to these resident macrophages in regulating cardiac lymphatic growth and patterning, along the base-apex axis and laterally to cover both dorsal and ventral surfaces of the developing heart. Tissue-resident macrophages were observed proximal and in direct contact with lymphatic vessels, where they accumulated at branch points. Moreover, in three independent mutant mouse models, *Pu.1^-/-^*, *Csf1r-CreER;R26R-DTA* and *Cx3cr1-CreER;R26R-DTA*, macrophage depletion resulted in a truncated and mis-patterned lymphatic plexus, suggesting an essential role in modulating the extension and branching of the lymphatic endothelium (Figs 3,5,6).

A previous study reported an analogous role for embryonic macrophages in the remodeling of the primitive coronary blood plexus (Leid et al., 2016). Here the authors reported that resident macrophages control the selective expansion of perfused blood vessels, and that an absence of macrophages resulted in hyper-branching of the developing coronaries (Leid et al., 2016). In contrast, we observed a transient reduction in vessel growth and branching (Fig. 4), and a mild, but reproducible mis-patterning of the coronary vessels following macrophage depletion in *Pu.1^-/-^*, *Csf1r-CreER;R26R-DTA* and *Cx3cr1-CreER;R26R-DTA* mutant mice (Figs. 4-6). Our findings are supported by studies documenting reduced vessel branching in alternative vascular beds of mice lacking embryonic macrophages, such as the developing hindbrain and postnatal retina (Fantin et al., 2010). Thus, the differences in severity of phenotype between our analyses and that of Leid and colleagues could be due to distinct mouse models used and/or approaches employed to characterize the coronary vasculature (whole-mount heart immunostaining in this study, versus analyses of cryosections in the previous (Leid et al., 2016)).

At a molecular level, the remodeling of the lymphatic vasculature could arise from macrophage phagocytic activity, release of soluble cytokines or cell-cell interactions. Tissue-resident macrophages have been reported to control various processes during vessel development, in a variety of blood and lymphatic vascular beds, mediated by secretion of trophic factors. These include, regression of the transient hyaloid endothelium by inducing WNT7B-driven cell death (Lobov et al., 2005), inhibition of branch formation in the developing lymphatic system in the diaphragm through secretion of an elusive factor (Ochsenbein et al., 2016), controlling LEC proliferation in dermal lymphatics (Gordon et al., 2010), mediating coronary blood plexus remodeling through selective expansion of the perfused vasculature *via* putative insulin-like growth factor (IGF) signaling (Leid et al., 2016). In contrast, our findings in the developing mouse heart and *in vitro* models of human lymphatic capillary tube formation and sprouting (Figs 3,5-7) favor a mechanism mediated by direct macrophage-endothelial cell physical contact/adhesion (Fig.1), similar to the role of macrophages in the developing hindbrain and retina vasculature (Fantin et al., 2010). In particular, a requirement for macrophage HA in lymphatic sprouting was identified (Fig. 7). HA is a key structural component of the vertebrate extracellular matrix and has recently been implicated in the postnatal development and response to adult injury (inflammation-driven) of murine corneal lymphangiogenesis (Sun et al., 2019), a process that also depends on the presence of macrophages (Maruyama et al., 2012). Hyaluronan levels are determined, partly by three hyaluronan synthases (HAS1-3), and three hyaluronidases (HYAL1-3) (Triggs-Raine & Natowicz, 2015). HA is also known to interact with different cell surface receptors, including LYVE-1, CD44 and HMMR/CD168 to directly influence endothelial cell motility, proliferation and survival, though the role of such receptors in the regulation of hyaluronan levels remains elusive (Savani et al., 2001; Triggs-Raine & Natowicz, 2015). As such, functional redundancy by different HA binding proteins likely contributes to the lack of a developmental (lymphatic) phenotype in *Lyve1* and *Cd44* single and compound knockout mice (Gale et al., 2007; Luong et al., 2009). Conversely, genetic disruption of hyaluronan synthesis abrogated normal cardiac morphogenesis leading to mid-gestation demise of mutant embryos and impaired the response of cardiac macrophages to ischemia reperfusion injury resulting in poor macrophage survival and functional cardiac output (Camenisch et al., 2000; Petz et al., 2019). Thus, further studies are needed to dissect out the HA/HA-binding receptor axis required for macrophage-induced lymphatic sprouting. Likewise, an indirect effect, *via* the release of a soluble growth factor(s) by tissue-resident macrophages cannot be categorically ruled out at this stage, albeit we can discount VEGF-C given the lack of expression by macrophages in the developing heart (Fig. S4).

We assign a novel function to the yolk sac-derived, tissue-resident macrophage lineage, against a back-drop of a rapidly evolving field which attributes ever increasing plasticity, in terms of cell fate and signaling, to these essential resident lineages. This is complemented by insights into our fundamental understanding as to how discrete organ-based vascular beds are formed and the degree of heterogeneity in terms of cellular contributions to organ-specific endothelium. Live imaging in the adult mouse and zebrafish recently demonstrated that inflammatory macrophages are required for orchestration of skin wound neoangiogenesis (Gurevich et al., 2018). Likewise, macrophages of diverse phenotypes supported vascularization of 3D human engineered tissues (Graney et al., 2020). Moreover, the maintenance of the mouse choroidal vascular network, as well as lymphatic vessels in the cornea is dependent on the presence of macrophages, with decreased macrophage number and activation leading to reduced lymphatic vessel formation and contributing to impaired diabetic wound healing in a mouse model of corneal wound healing (Maruyama et al., 2007; Maruyama et al., 2012; Yang et al., 2020). Importantly the close interaction between cardiac lymphatics and macrophages during development, appears to manifest in the postnatal heart via infiltrating inflammatory macrophages and injury-activated lymphangiogenesis following myocardial infarction (Klotz et al., 2015; Vieira et al., 2018). The functional role for macrophages in directly regulating lymphatic growth and remodeling during development we report herein may, therefore, manifest in response to injury in the adult setting. Taken together, this further strengthens the rationale for extrapolating from our developmental studies to therapeutically target macrophage-lymphatic responses during postnatal cardiovascular disease/injury.

## MATERIAL AND METHODS

Key resources table

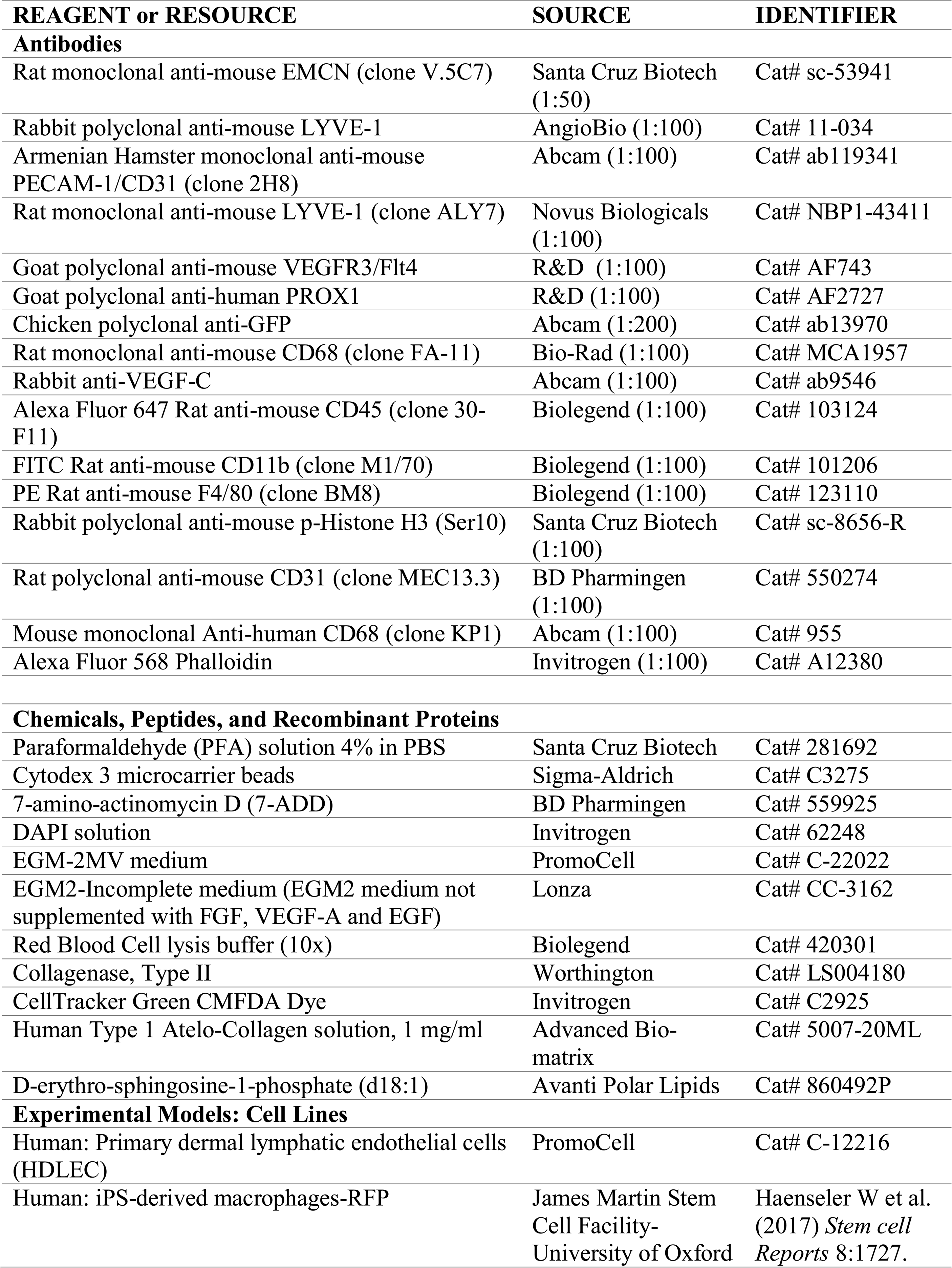
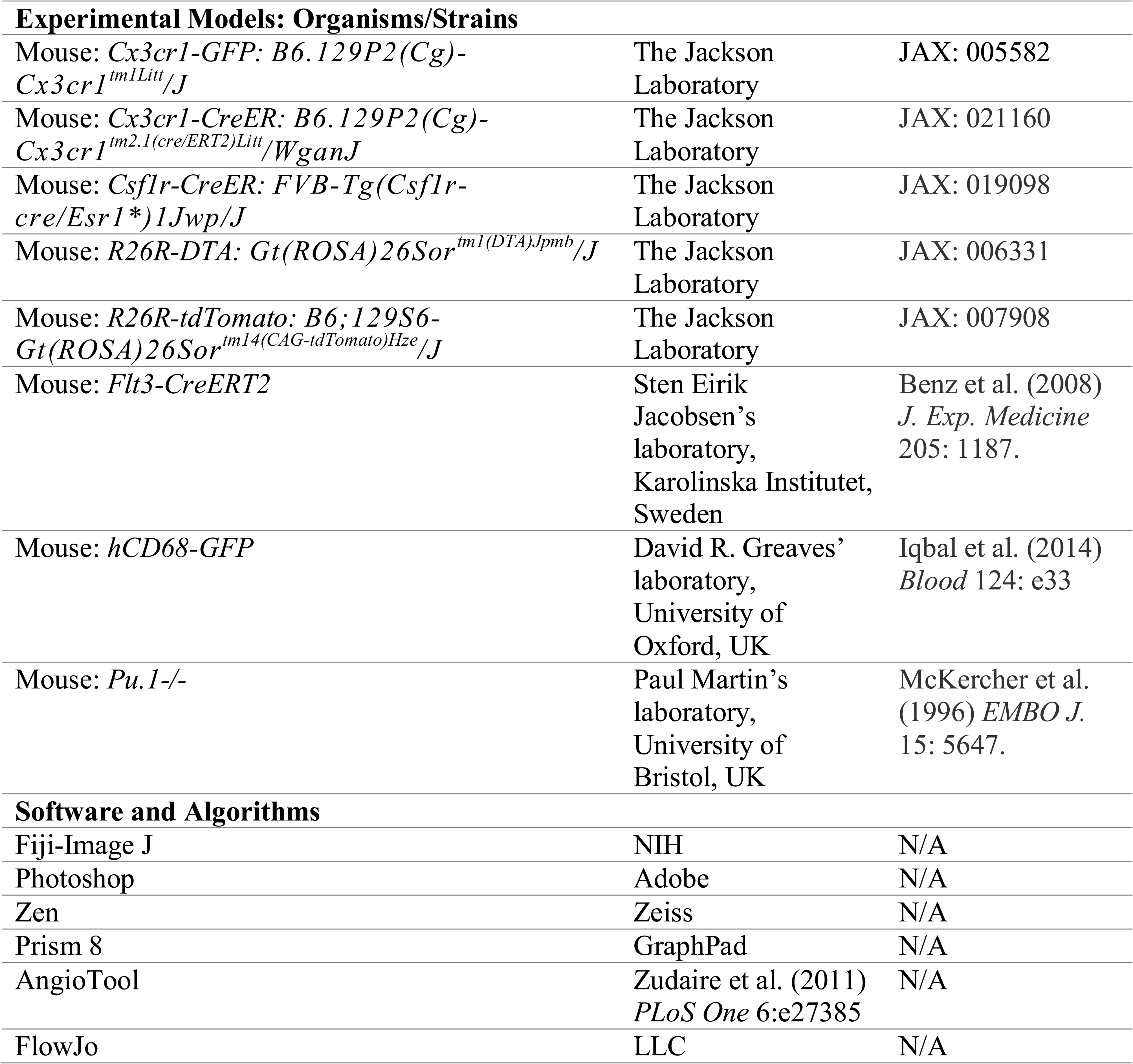

### Mouse lines

Genetically modified mouse lines used in the study were kept in a pure C57BL/6 genetic background and are listed in **Key resources table**. *Cx3cr1-GFP* knock-in mice and *hCD68-GFP* transgenic mice were crossed with C57BL/6 to generate *Cx3cr1^GFP/+^* and *hCD68-GFP^+^*samples, respectively. *Pu.1^+/-^* mice were intercrossed to generate *Pu.1^-/-^* specimens. *Csf1r-CreER* transgenic mice, *Cx3cr1-CreER* knock-in mice and *Flt3-CreERT2* transgenic mice were crossed with *R26R-tdTomato* or *R26R-DTA* reporters strains to generate *Csf1r-CreER;tdTomato*, *Csf1r-CreER;R26R-DTA*, *Cx3cr1-CreER;R26R-tdTomato*, *Cx3cr1-CreER;R26R-DTA* and *Flt3-CreERT2;tdTomato* mice. *Pu.1^-/-^;Csf1r-CreER;tdTomato* compound transgenic mice were generated by crossing *Pu.1^+/-^;Csf1r-CreER* with *Pu.1^+/-^ ;R26R-tdTomato* mice. Both males and females were used in the study. For timed-mating experiments, 8-16-week-old mice were set up overnight and females checked for vaginal plugs the following morning; the date of a vaginal plug was set as embryonic day (E) 0.5. For tamoxifen-dependent tissue-specific gene activation, 75mg/Kg/dose of tamoxifen (Sigma) were administered to pregnant dams, intraperitoneally (i.p.). Mice were housed and maintained in a controlled environment by the University of Oxford Biomedical Services. All animal experiments were carried out according to the UK Home Office project license (PPL) 30/3155 and PPDE89C84 compliant with the UK animals (Scientific Procedures) Act 1986 and approved by the local Biological Services Ethical Review Process.

### Cell lines

Primary human dermal lymphatic endothelial cells (HDLEC) isolated from the dermis of juvenile foreskin (PromoCell) were cultured in presence of endothelial cell growth medium (ECGM)-MV2 (PromoCell), according to the manufacturer’ instructions. Human iPS-derived macrophages constitutively expressing RFP (Haenseler et al., 2017) were obtained from the James Martin Stem Cell Facility (Sir William Dunn School of Pathology, University of Oxford, UK). The human iPS cell line was derived previously from dermal fibroblasts of a healthy donor that had given signed informed consent for the derivation of iPS cells (Ethics Committee: National Health Service, Health Research Authority, NRES Committee South Central, Berkshire, UK (REC 10/H0505/71)). iPS-macrophages were cultured in macrophage medium composed of X-VIVO™15 (Lonza), GlutaMax (Invitrogen) and M-CSF (Invitrogen), as previously described (van Wilgenburg et al., 2013). Cell lines were maintained in a humidified atmosphere of 5% CO_2_ at 37°C.

### Sample preparation for immunostaining

Embryos were harvested at the required embryonic stage, placed in ice-cold PBS (Sigma) and the amniotic sac was removed. The heart was micro-dissected from the embryo for immunostaining experiments using fine forceps. Dissected hearts were fixed for 6 hours in 2% PFA/PBS at room temperature, permeabilized in 0.3% Triton X-100/PBS (twice, 10 minutes each) and then blocked in 1% BSA, 0.3% Triton X-100 in PBS for at least 2 hours. Samples from fluorescent reporter lines (e.g. *Cx3cr1-GFP* and *CD68-GFP*) were protected from light throughout this procedure. Samples were incubated with primary antibodies (diluted in block; listed in **Key resources table**) overnight at 4°C, then washed three times for at least 1 hour each in 0.3% Triton X-100/PBS. Samples were incubated with Alexa Fluor®-conjugated secondary antibodies (diluted in PBS; Invitrogen) overnight at 4°C, protected from light, then rinsed three times for at least 10 minutes each with PBS. The hearts were then orientated according to dorsal-ventral surface aspect and mounted in 50% glycerol/PBS in glass-bottomed dishes (Mattek). Imaging was performed using an Olympus FV 1000 or Zeiss LSM 780 scanning confocal microscope. Images were digitally captured and processed using Zen and FIJI-Image J software. Analysis of total vessel length and junction number calculation were performed using AngioTool software (Zudaire et al., 2011).

### Preparation of single cell suspensions from the heart

Fetal hearts harvested for flow cytometry studies were isolated, placed in ice-cold HBSS (Life Technologies), finely minced into small pieces, and digested with collagenase type II (Worthington Laboratories) solution (containing 500units/ml HBSS) at 37°C for 30 minutes with agitation at 180 rpm. Dissociated samples were then passed into a 50-mL conical tube (Corning) through a 70-μm cell strainer, then rinsed with 3 mL ice-cold HBSS and transferred back to a 15-mL conical tube. Samples were spun for 7 minutes at 350 *g* at 4°C, and the supernatant carefully discarded. Cell pellets were resuspended in 5 mL of Red Blood Cell (RBC) lysis buffer (BioLegend) and incubated at room temperature for 10 minutes, followed by a repeat centrifugation at 350 *g* at 4°C for 7 minutes. The RBC lysis buffer was removed, and cells resuspended in 2% FBS/PBS. Isolated single cardiac cells were stained and subjected to flow cytometry analyses.

### Flow cytometry

Using 7-AAD exclusion (BD Pharmingen), only live cells were analysed. All antibodies used for flow cytometry are listed in **Key resources table**. Flow cytometric analyses were performed using FACSAria III flow cytometer (BD Biosciences) and FlowJo software (LLC). Samples resuspended in 2% FBS/PBS solution were blocked with mouse Fc Block (Miltenyi Biotec) for 5 minutes on ice, followed by labelling for 20 minutes at room temperature with each antibody combination.

### Time-lapse tube formation assay

384-well plates (PerkinElmer) were coated with Growth Factor-reduced Matrigel (Corning) and incubated for 30 minutes at 37°C. 25 μL of ECGM-MV2 containing 4,500 HDLECs were added per well and incubated for 60 minutes at 37°C. Then, 25 μL of macrophage media containing 2,250 human iPS-derived macrophages-RFP were added per well. For time-lapse imaging acquisition, an Evos® FL Auto Cell Imaging System (ThermoFisher) was used and one image was taken every 8 minutes for 15 hours. At the end of image acquisition, cells were fixed with 4% PFA solution (Santa Cruz Biotech) for 30 min and washed with PBS for further analysis (e.g. immunostaining).

### Immunostaining of HDLEC-macrophage co-cultures

HDLEC-like tubes were permeabilized with 0.1% Triton X-100/PBS and then incubated with a staining solution containing Alexa Fluor® 568 Phalloidin and DAPI (both Invitrogen). Immunofluorescence staining was performed using an antibody against human CD68 (Abcam).

### iPS-derived macrophage transfection

Macrophages were transfected in a 24-well plate as previously described (Troegeler et al., 2014). Macrophages were washed twice with warm complete XVIVO in order to remove floating cells. 250 μL of fresh complete XVIVO were added per well and cells kept at 37LJC and 5% CO2. 3,75 uL of siRNA (20 uM - Thermofisher), 110.25 uL XVIVO depleted (without MCSF) and 11 uL HiPerfect (Qiagen) were gently mixed in a tube and incubated at room temperature for 15 to 20 min. 125 uL of the mix was then added in each well drop by drop. The plate was gently rocked to assure nice distribution of the mix. After 6 hours, 0.5 mL of Complete XVIVO was added and the cells were harvested 3 days later for RNA extraction and/or experimental procedures.

### HAase treatment and immunostaining

Macrophages were seeded on coverslips and treated with 15 U/mL or 30 U/mL hyaluronidase (HAase - Sigma) 2 hours prior experiment. For staining, cells were fixed with 4% PFA and permeabilized in 0.2% Triton X-100/PBS for 10 min and then blocked in 1% BSA, 0.1% Triton X-100 in PBS for at least 2 hours. Macrophages were incubated with 3 ug/mL biotinylated Hyaluronan binding protein (Amsbio) overnight at 4°C, then washed three times with PBS. Samples were incubated with Alexa Fluor® 488 Streptavidin (diluted in PBS; Biolegend) overnight at 4°C, protected from light, then rinsed three times for at least 10 minutes each with PBS. Coverslip were then mounted on slides using Vectashield + DAPI (Vector Laboratories).

### Microbeads capillary sprouting assay

The capillary sprouting assay was performed as previously described (Schulz et al., 2012). HDLECs were incubated in a 15-mL conical tube in presence of cytodex 3 beads (Sigma) at a ratio of 400 cells per bead in EGM-2MV medium (Lonza) for 4 hours at 37°C with shaking every 20 minutes. Beads were then incubated in a cell flask at 37°C for 48 hours in EGM-2MV medium. The HDLEC-coated beads were subsequently collected and labelled with 2 μM Cell Tracker Green dye (Invitrogen) for 30 minutes, embedded in 1mg/mL Type I collagen hydrogel (Advanced Bio-Matrix), containing 2 μM D-erythro-sphingosine-1-phosphate (Avanti Polar Lipids), and cultured in black clear-bottom 96-well plates (Perkin-Elmer) in the presence or absence of 10,000 human iPS-derived macrophages-RFP per well. After 60 minutes of incubation, a 1:1 mix containing EGM2-Incomplete (EGM2 media without supplementation with EGF, FGF2 and VEGF-A; Lonza) and macrophage media (v/v) was added and the beads incubated for two days at 37°C. Then, the beads were fixed with 4% PFA for 30 minutes and washed twice with PBS. Each well was imaged with a Zeiss LSM 780 scanning confocal microscope and the number of sprouts per bead calculated.

### Spheroids sprouting assay

Spheroids of 400 HDLECs were generated using a 24-well plate Aggrewell 400 (Stemcells Technologies) using EGM2-Incomplete. The following day, spheroids were collected and embedded in 1mg/mL Type I collagen hydrogel and cultured in black clear-bottom 384-well plates (Perkin-Elmer) in the presence or absence of 2,5k human iPS-derived macrophages-RFP per well. After 60 minutes of incubation, a 1:1 mix containing EGM2-Incomplete (Lonza) and macrophage media (v/v) was added and the spheroids incubated for two days at 37°C. Then, the spheroids were fixed with 2% PFA for 30 minutes, washed twice with PBS and stained with Alexa Fluor® 488 phalloidin (Invitrogen). Each well was imaged with an automated cell imaging system (Pico microscope - Molecular device) and the number of sprouts per spheroids calculated.

### RNA extraction and qRT-PCR

Total RNA was extracted using Trizol Reagent (Invitrogen). Reverse transcription (RT) was performed using SuperScript(R) III Reverse Transcriptase (Invitrogen), as recommended by the manufacturer. Primer sequences (Invitrogen) used were: *36B4*_F CTACAACCCTGAAGAAGTGCTTG *36B4*_R CAATCTGCAGACAGACACTGG *CSF1R*_F CCTCACTGGACCCTGTACTC *CSF1R*_R GGAAGGTAGCGTTGTTGGTG *CX3CR1*_F ACCAACTCTTGCAGGTCTC *CX3CR1*_R TGTCAGCACCACAACTTGG *ITGB2/CD18*_F GTCTTCCTGGATCACAACGC *ITGB2/CD18*_R CAAACGACTGCTCCTGGATG *ITGAM/CD11b*_F CAGTGAGAAATCCCGCCAAG *ITGAM/CD11b*_R CCGAAGCTGGTTCTGAATGG *VEGFC*_F GGAGGCTGGCAACATAACAG *VEGFC*_R TTTGTCGCGACTCCAAACTC *LYVE1*_F CTGGGTTGGAGATGGATTCG *LYVE1*_R TCAGGACACCCACCCCATTT *CD44_*F AAGTGGACTCAACGGAGAGG *CD44_*R GTCCACATTCTGCAGGTTCC *HMMR/CD168*_F GGCTAAGCAAGAAGGCATGG *HMMR/CD168*_R TCCCTCCAGTTGGGCTATTT RealLJtime polymerase chain reaction (PCR) assays were run on a ViiA 7 step realLJtime PCR machine (AppliedLJBiosystems). Normalization was performed using *36B4* as a reference gene. Quantification was performed using the comparative Ct method.

### scRNA-seq and analysis

E7.75_E8.25_E9.25 10x Chromium data (GSE126128) were downloaded from UCSC Cell Browser as raw UMI count matrix (de Soysa et al., 2019). Only the E9.25 datasets were selected for further analysis. E10.5 heart 10x Chromium data were downloaded as raw counts from GEO (GSE131181) (Hill et al., 2019). All scRNA-seq datasets were analyzed using Seurat (Butler et al., 2018; Stuart et al., 2019) in R as follows: individual replicates were integrated using SCTransform method, principal component analysis was used to define the cell clusters, which were visualized with the UMAP method (Becht et al., 2018; Butler et al., 2018).

### Statistical analysis

All data are presented as mean ± standard error of the mean (SEM). Statistical analysis was performed on GraphPad Prism 8 software. The statistical significance between two groups was determined using an unpaired two-tailed Student’s *t*-test, these included an F-test to confirm the two groups had equal variances. Among three or more groups (e.g. data shown in Fig. 1A), one-way analysis of variance (ANOVA) followed up by Tukey’s multiple comparison test was used for comparisons. A value of p ≤ 0.05 was considered statistically significant.

### Contact for reagent and resource sharing

Further information and requests for resources and reagents should be directed to and will be fulfilled by the Lead Contact, Paul R. Riley.

## Acknowledgements

We would like to thank Paul Martin (School of Biochemistry, University of Bristol, Bristol, UK) for the kind permission to use the *Pu.1^-/-^* mice on a C57BL/6 genetic background; Micron Oxford Advanced Bioimaging Unit for access to and training in the use of confocal microscopy; and the staff of the University of Oxford Biomedical Services Building for excellent mouse husbandry. We thank the staff at the James Martin Stem Cell Facility, which has received core support from the Oxford Martin School (LC0910-004), Wellcome Trust (WTISSF121302) and MRC (MC EX MR/N50192X/1) for assistance with iPS-macrophage differentiation. This work was funded by the British Heart Foundation (chair award CH/11/1/28798 and programme grant RG/08/003/25264 to PRR) and supported by the BHF Oxbridge Centre of Regenerative Medicine (RM/13/3/30159); a Wellcome Trust Doctoral Training Fellowship 106334/Z/14/Z to TJC; a Wellcome Trust Four year PhD Studentship 215103/Z/18/Z to KK; a BHF Intermediate Basic Science Research Fellowship FS/19/31/34158 to JMV; a British Israel Research and Academic Exchange Partnership (BIRAX) Grant 13BX14PRET; a Leducq Foundation Transatlantic Network of Excellence Program 14CVD04 and MRC Unit funding to DGJ.

## Competing interests

PRR is co-founder and equity holder in OxStem Cardio, an Oxford University spin-out that seeks to exploit therapeutic strategies stimulating endogenous repair in cardiovascular regenerative medicine.

## Authors Contributions

TJC, XS, JMV and PRR conceived and designed the study. TJC, XS, CVDC and JMV carried out all mouse experiments, including sample generation, immunostaining and imaging. CR performed the macrophage-lymphatic endothelial cell co-culture experiments. TJC performed the flow cytometry studies and analyzed the data generated. IEL performed scRNA-seq analysis and generated UMAP and dot plots. AML and SEWJ generated the *Flt3-CreERT2;tdTomato* samples. DRG provided the *hCD68-GFP* reporter mice. DGJ provided significant advice and technical assistance with macrophage-lymphatic cell interactions and involvement of the macrophage hyaluronan surface glycocalyx. CB, SAC and WJ provided iPS-derived macrophages-RFP and technical assistance with co-culture studies. RPC and PRR co-supervised TJC. XS, TJC, CR, KK and JMV analyzed the data. JMV wrote the manuscript. TJC, XS, JMV and PRR edited the manuscript. JMV and PRR supervised the study.

---

**Fig. S1. Tissue-resident macrophages are present in the developing murine heart at E10.5.** (A) UMAP plot demonstrating the different major clusters in the E10.5 heart scRNA-seq dataset (total 12,535 cells, *n* = 3 batches). (B) Dot plot showing proportion of cells in each cluster expressing selected genes. Dot size represents percentage of cells expressing, and color scale indicates average expression level. (C-E) Whole-mount immunostaining for GFP (green), and PECAM-1 (red) to visualize tissue-resident macrophages in the sinus venosus and ventricular surface of hearts-derived from *Cx3cr1^GFP/+^* embryos at E10.5. AvCu, atrioventricular cushion; CM_A, atrial cardiomyocytes; CM_AVC, atrioventricular canal cardiomyocytes; CM_OFT, cardiomyocytes of outflow tract; CM_Prol, proliferative cardiomyocytes; CM_S, septal cardiomyocytes; CM_SV, sinus venosus cardiomyocytes; CM_Trab, trabecular cardiomyocytes; CM_V, ventricular cardiomyocytes; Endo, endocardial cells; EndoMT, endocardial epithelial-to-mesenchymal transition; Epi, epithelial cells; Mes, mesenchyme; RBC, red blood cells; SHF, second heart field. Scale bar 100 μm

**Fig. S2. Loss of macrophages is not associated with hyperproliferation of the cardiac lymphatic endothelium.** (A,B) Representative flow cytometry histograms of cells isolated from control (Co) and *Pu.1^-/-^* hearts at E14.5 labeled against CD45 and CD11b (A) or F4/80 and CD11b markers (B). (C,D) Quantification of the myeloid (C), defined as CD45^+^CD11b^+^, and macrophage populations, defined as CD45^+^CD11b^+^F4/80^+^ (D) residing in Co versus *Pu.1^-/-^* hearts at E14.5. Data represent mean ± SEM; *n* = 4 hearts per group. Significant differences (*p* values) were calculated using an unpaired, two-tailed Student’s *t*-test (\*\**p* ≤ 0.01). (E,F) Histological characterization of control and *Pu.1^-/-^* hearts at embryonic day (E) 16.5 using Hematoxylin & Eosin staining. (G-N) Whole-mount immunofluorescent staining for phospho-histone H3 (pH3; green) and PROX1 (red) in control (Co; G-J) and *Pu.1^-/-^* hearts (K-N) at E16.5. All scale bars 100 μm, except E 1mm.

**Fig. S3. Human induced pluripotent stem cell (hiPS)-derived macrophages display HA on their cell surface, express HA-binding proteins and adhesion receptor CD18/CD11b but lack the expression of *VEGFC*.** (A) Marker expression analysis by qRT-PCR using RNA isolated from hiPS-macrophages. Data presented as mean ± SEM; *n* = 6 independent experiments, with three replicates/experiment. (B-D) Representative HA staining (green) on hiPSC-derived macrophages-RFP (red) under control conditions (B) or following incubation with HAase 15 U/mL (C) and HAase 30 U/mL (D). DAPI (blue) labels the nuclear DNA. Note the loss of cell surface (glycocalyx)-associated HA following treatment with HAase 30 U/mL. (E) *ITGB2* and (F) *ITGAM* expression analysis by qRT-PCR using RNA isolated from control hiPS-macrophages (macrophages) or hiPS-macrophages treated with control siRNA or siRNA oligonucleotides against *ITGB2*/*CD18* (two independent siRNA sequences were used; siCD18 #1 and siCD18 #2). Data presented as mean ± SEM; *n* = 6 independent experiments, with three replicates/experiment. Significant differences (*p* values) were calculated using one-way ANOVA followed up by the Tukey’s multiple comparison test (\*\*\*\**p* ≤ 0.0001). All scale bars 200 μm.

**Fig. S4. Tissue-resident macrophages in the developing mouse heart display hyaluronan on their cell surface and lack expression of VEGF-C.** (A-F) Whole-hearts from *Cx3cr1-GFP* embryos were analyzed for GFP (green), LYVE-1 (red) and VEGF-C (white) expression at E16.5. (B-F) Magnified views of box shown in (A). Note VEGF-C co-localization with LYVE-1 on lymphatic endothelium, but not on LYVE-1 and GFP co-expressing macrophages. (G-J) GFP (green) and VEGF-C (red) immunostaining of tissue sections derived from E16.5 hearts documenting lack of expression of VEGF-C in CX3CR1-GFP+ ventricular/subepicardial macrophages (G,H) and enrichment for VEGF-C in the coronary stems (white arrows; I, J). (H,J) Magnified views of boxes shown in (G,I). (K-P) Whole-hearts from *Cx3cr1-GFP* embryos were analyzed for GFP (green), HA (red) and LYVE-1 (white) expression at E16.5. (L-P) Magnified views of box shown in (K). Note HA colocalization with GFP and LYVE-1 in tissue-resident macrophages located in the subepicardial space, contiguous to the expanding lymphatic vasculature (white arrowheads in L). (Q-T) GFP (green), HA (red) and LYVE-1 (white) immunostaining of tissue sections derived from E16.5 hearts indicating expression of HA in CX3CR1-GFP+ subepicardial macrophages (white arrowheads in Q). DAPI (blue) labels nuclear DNA. LV, left ventricle; OFT, outflow tract. All scale bars100 μm, except H 25 μm and J,Q 50 m.

**Movie S1.** Time-lapse imaging of tube formation in co-cultures of human primary lymphatic endothelial cells (phase contrast) and human-iPS-derived macrophages expressing RFP.

